# Amyloid stability and condensate material properties independently govern the hnRNPA2 condensate to fibril transition

**DOI:** 10.64898/2026.11.04.764024

**Authors:** Daniel J. Kieffer, June Ho Hwang, Michael K. Rosen, Lukasz A. Joachimiak

## Abstract

hnRNPA2 is a component of cellular membraneless organelles many of which contain RNA and RNA-binding proteins (RBPs). RBPs frequently contain low-complexity domains (LCDs) that can also assemble into amyloid fibrils. Fibrils isolated from patients with frontotemporal lobar degeneration contain ordered segments of these LCDs and disease-associated mutations can enhance the homotypic interactions that promote aggregation. Although previous studies have largely examined the condensate-to-fibril transition by altering condensate stability, how amyloid structure governs this process remains poorly understood. Here, we show that the hnRNPA2 LCD forms a reproducible fibril fold and by combining structure-guided thermodynamic profiling, peptide design, mutagenesis and in vitro characterization under conditions with (two-phase) and without (one-phase) condensates, we define the molecular determinants of its assembly. We identify an aromatic triad within the fibril core that controls fibril formation kinetics and thermodynamic stability. Mutational analysis further reveals that condensate stability and fibril stability are largely independent. Finally, microrheology linked reduced condensate viscosity to faster fibril formation, supporting a role for condensate material properties in regulating aggregation kinetics. Together, these findings support a model in which the condensate-to-fibril transition is governed by the interplay between a disordered “fuzzy coat” and a fibril-forming aromatic triad within hnRNPA2. This work provides a structural framework for understanding how condensates can transition into amyloid assemblies.

**Significance Statement:** Protein condensates may promote or suppress amyloid formation, but the properties governing these outcomes remain poorly understood. Through mutational analysis, this study examines the relationships among condensate material properties, condensate and fibril stability, and aggregation kinetics. The findings provide a framework for understanding how changes in condensate properties influence amyloid formation in the system studied, with potential implications for disease-associated protein assembly.

## Introduction

Biomolecular condensates are membraneless organelles that form through liquid-liquid phase separation (LLPS). LLPS is the prevailing model for the formation of dynamic biological structures like cytoplasmic stress granules[1, 2]. These membraneless compartments form through networks of homotypic and heterotypic interactions among RNA molecules, and RNA-binding proteins (RBPs), including TDP-43, FUS and hnRNPA2[2–6]. These RBPs frequently contain low-complexity domains (LCDs) which drive condensation[6, 7]. RBPs are associated with neurodegenerative diseases, including amyotrophic lateral sclerosis (ALS), frontotemporal dementia (FTD) and multisystem proteinopathy (MSP)[8–10] [11]. In these disorders, RBPs such as TDP-43, TAF15, hnRNPA2 and hnRNPA1 can mislocalize from the nucleus to the cytoplasm, and through a phase separated intermediate undergo pathological conversion into β-sheet-rich aggregates[9, 10, 12]. These assemblies are defining pathological hallmarks of several RBP proteinopathies and are thought to contribute to cellular dysfunction and neurodegeneration[11, 13–19].

Disease-associated mutations often reside within the low-complexity domains (LCDs) of RBPs, which are intrinsically disordered regions enriched in repetitive, flexible, aromatic, polar and charged residues. These LCDs can drive assembly into biomolecular condensates through multivalent, weakly adhesive interactions, including aromatic, electrostatic, cation-π and π-π contacts[20, 21]. The same sequence features can also promote amyloid formation by enhancing homotypic interactions and β-sheet assembly[6]. This overlap has led to the hypothesis that concentrating RBPs within condensates, especially at condensate interfaces[22–25], promotes intra-and intermolecular contacts needed for fibril formation. However, recent work suggests that the dynamic interaction network within condensates can be inhibitory to amyloid formation by sequestering soluble protein which prevents conversion into β-sheet-rich amyloids[26]. New methods now enable quantification of condensate material properties[27, 28]. These have revealed that increased viscoelasticity can influence the balance between condensation and amyloid formation, in some cases suppressing amyloid assembly[29]. Thus, a key gap remains in understanding how the interactions that drive condensate formation are related to RBP amyloid formation.

While most work examining this relationship has focused on how these interactions alter the condensate phase, few studies have examined this relationship in the context of amyloid structure. Recent advances in cryo-EM have enabled high-resolution structures of amyloid fibrils, including those derived from patient material, providing new insight into how fibril architecture relates to disease. However, in vitro amyloid assembly remains difficult to control for many proteins, often producing structural polymorphism that complicates mechanistic interpretation[30]. hnRNPA2 is somewhat unusual in forming a monomorphic fibril structure[31], making it a powerful system for linking sequence features, condensate behavior and amyloid architecture. In this structure, the ordered fibril core is flanked by an aromatic-rich fuzzy coat formed by N-and C-terminal low-complexity sequences. These sequences extend beyond the ordered amyloid core to form a fuzzy coat. How this region contributes to condensate formation and influences fibril assembly remains incompletely understood[35]. This architecture defined by cryo-EM is also consistent with prior molecular-footprinting studies showing that buried and solvent-accessible aromatic residues can be correctly distinguished by this approach[32]. In addition, hydrogel partitioning experiments identified key residues required for hnRNPA2 templating[32]. Together, these features make hnRNPA2 an excellent model for testing how interactions that drive phase separation are connected to amyloid formation from a structural perspective.

Here we combined cryo-EM of in vitro amyloids, in silico thermodynamic profiling, peptide design, phase separation assays, aggregation measurements and video particle tracking microrheology (VPT) to define how hnRNPA2 transitions from a dynamic condensate to an ordered amyloid state. First, cryo-EM revealed a defined hnRNPA2 fibril fold formed in vitro. Within individual fibrils, we also observed a structural variant that preserves the core fold but adopts distinct conformations at the N- and C-termini. These experiments identified an aromatic triad, centered on Y275, Y283 and F309, as a major stabilizing element of the hnRNPA2 fibril fold. Mutational LCD and peptide fragment-based analyses showed that the aromatic network promotes fibril formation in both the one-phase regime, where monomer converts directly to fibril and the two-phase regime, where monomer first partitions into condensates before undergoing amyloid conversion. By contrast, fuzzy-coat mutations enhanced fibril formation under two-phase conditions while retaining one-phase aggregation half-times broadly comparable to WT. Finally, we used microrheology to measure hnRNPA2 condensate material properties and found that mutations within the fuzzy coat decrease droplet viscosity and are associated with faster fibril formation. Together, these data indicate that the condensate-to-fibril transition is governed by two coupled factors: sequence-encoded interactions that stabilize the amyloid structure and condensate material properties that regulate access to aggregation-competent states.

## Results

### The hnRNPA2 LCD adopts a defined and reproducible amyloid conformation

Previous work demonstrated that the low-complexity domain (LCD) of hnRNPA2 forms a highly reproducible amyloid structure, an unusual feature given the structural polymorphism commonly observed for fibrils assembled in vitro [31, 33]. To establish a structural framework for defining the interactions that stabilize this fold, we first reproduced the hnRNPA2 LCD fibril using an mCherry-hnRNPA2-LCD construct. Protein was purified as described previously [31, 33] and assembled into fibrils under quiescent conditions (SI Appendix, Fig. 1A). Cryo-EM analysis yielded a 1.99 Å reconstruction of the hnRNPA2 LCD fibril core spanning residues 263-319, with a helical twist of −2.99° and rise of 4.78 Å (Fig. 1A and SI Appendix, Fig. 1B-D, Table 1). These parameters closely match the previously reported structure and direct alignment of the two models gave a Cα RMSD of 0.375 Å (Fig. 1B) [31]. We additionally resolved a second fibril population at 2.12 Å resolution with a slightly different helical twist (−2.71°) and rise (4.80 Å) (SI Appendix, Fig. 2A-B, Table 2). Particles that resolve this second fibril population are within the same fibril as the first fibril population (SI Appendix, Fig. 2A). The ordered core of this structure spans residues 275-321 and closely superimposes on the central region of the main fibril conformation, with a Cα RMSD of 0.168 Å across residues 275-309 (SI Appendix, Fig. 2C-D). Differences are localized near the N- and C-terminal edges of the core, where residues extending through the PY-NLS motif[34, 35] adopt an alternative path around the fibril and generate distinct interlayer contacts (SI Appendix, Fig. 2E-F). Additionally, the N-terminal edge lacks residues G263-G274 suggesting structural flexibility at both the N and C-term edges. Thus, hnRNPA2 shows limited heterogeneity at the edge of the ordered core while preserving a highly reproducible central amyloid architecture.

**Figure 1.**
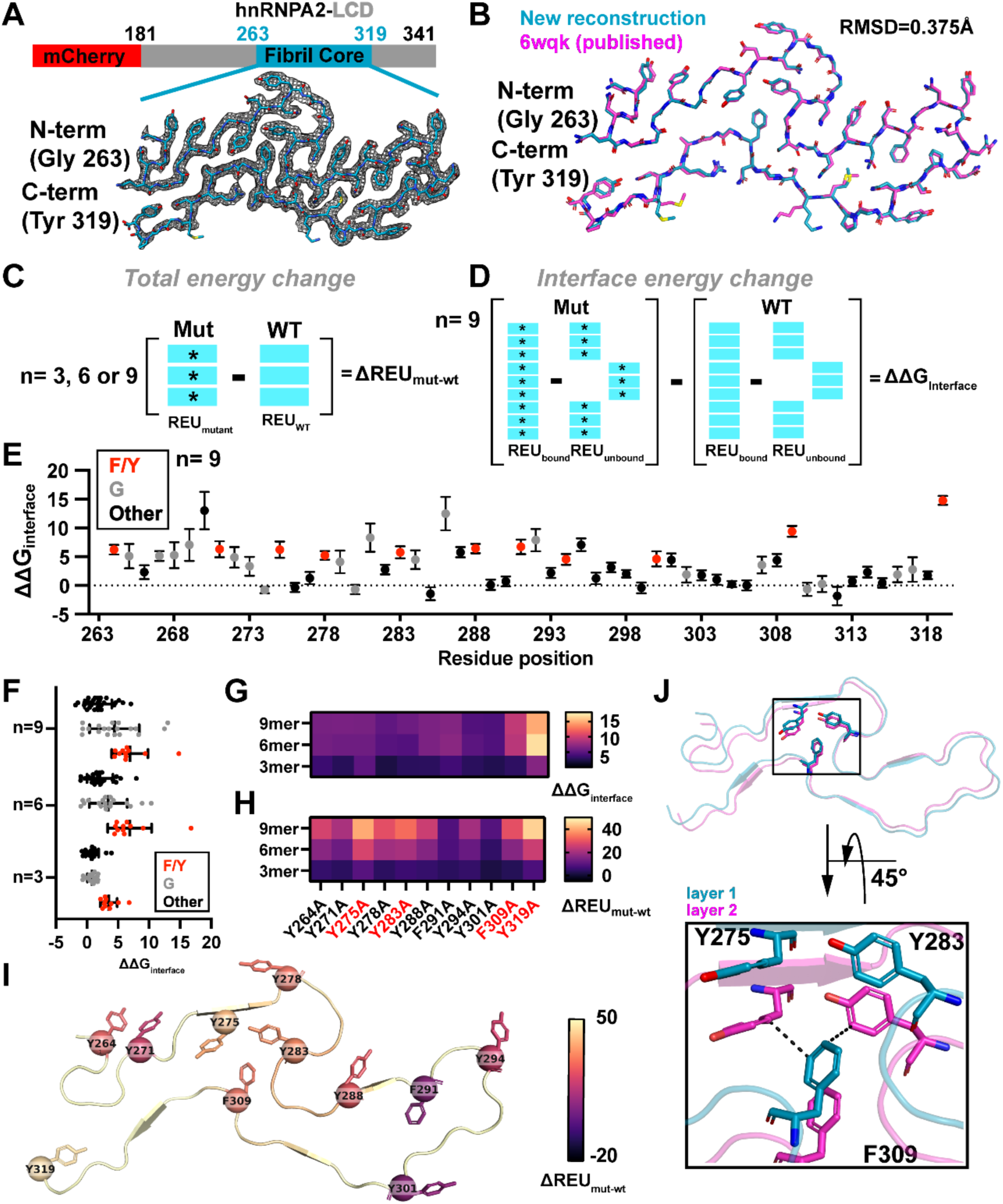
hnRNPA2 aromatic triad (Y275, Y283, F309) stabilizes the fibril fold. **(A)** Domain organization of the mCherry-hnRNPA2-LCD construct and cryo-EM density for the ordered fibril core spanning residues G263-Y319. **(B)** Structural alignment of the new hnRNPA2 fibril reconstruction with the previously reported structure (PDB 6WQK), yielding a Cα RMSD of 0.375 Å. **(C)** Schematic of the computational alanine-scanning strategy used to calculate the total energetic effect of mutation (ΔREU_mut-WT_) in 3-, 6- and 9-layer fibril assemblies. **(D)** Schematic of the interface-energy calculation used to determine ΔΔG_interface_ by comparing bound and unbound WT and mutant states. **(E)** Residue-by-residue ΔΔG_interface_ values for the 9-layer fibril assembly. Aromatic residues (Phe/Tyr), glycines and other residues are indicated separately. Data are shown as an average across 35 replicates with error bars representing standard deviation. **(F)** Distribution of ΔΔG_interface_ values for alanine substitutions in 3-, 6- and 9-layer fibril assemblies, grouped as aromatic residues, glycines, or other residues. Points represent mean values for each position. Data are shown as averages across F/Y, G and other group members with errors shown as standard deviation. **(G)** Heatmap of interface-energy changes for aromatic-to-alanine substitutions across 3-, 6- and 9-layer assemblies. Data are shown as averages across 35 replicates. **(H)** Heatmap of total energetic changes (ΔREU_mut-WT_) for aromatic-to-alanine substitutions across the three assembly sizes. Y275, Y283, F309 and Y319 are highlighted. Data are shown as averages across 35 replicates. **(I)** Structural mapping of aromatic residues onto the hnRNPA2 fibril core, colored according to the calculated energetic effect of alanine substitution. **(J)** Structural view of the Y275/Y283/F309 aromatic network across adjacent fibril layers. The enlarged view illustrates interlayer packing of F309 with Y275 and Y283.

**Figure 2.**
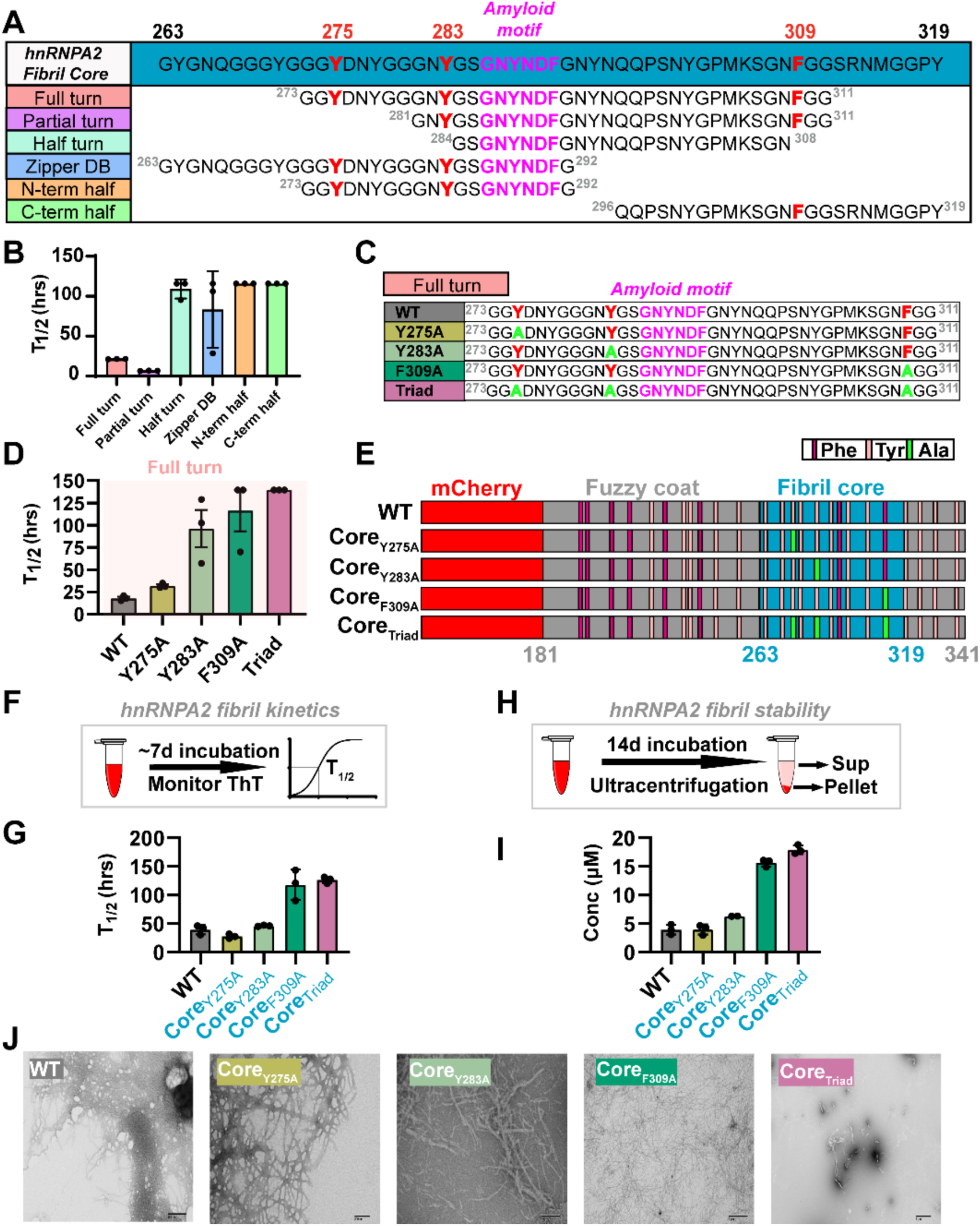
The aromatic triad is necessary for fibril formation. **(A)** Peptide constructs spanning different regions of the hnRNPA2 fibril core used to define sequence requirements for amyloid formation. The full-turn peptide spans residues 273-311 and contains Y275, Y283 and F309; progressively truncated peptides retain different portions of the fibril turn and aromatic network. ZipperDB-derived segment and N- and C-terminal half peptides are also shown. Amyloid motif is shown in pink. Aromatic triad (Y275, Y283 and F309) is indicated in red. **(B)** Aggregation half-times (T_1/2_) for the indicated fibril-core peptides. Peptides containing both Y283 and F309 are efficient in fibril formation. Data are shown as an average across three technical replicates with errors represented as standard deviation. Peptides that failed to reach measurable aggregation are plotted at the assay limit. **(C)** Schematic of full turn peptide constructs. Core aromatics labeled as red, alanine substitutions as green and amyloid motif as purple. **(D)** Aggregation half-times (T_1/2_) for the indicated fibril-core peptides. Y275 is dispensable for fibril formation, whereas Y283A, F309A and Triad substitutions impair fibril formation. Data are shown as an average across three technical replicates with errors represented as standard deviation. Peptides that failed to reach measurable aggregation are plotted at the assay limit. **(E)** Schematic of the mCherry-hnRNPA2-LCD constructs used for LCD fibril experiments. Individual Y275A, Y283A and F309A substitutions and the combined Y275A/Y283A/F309A mutant _(CoreTriad)_ are indicated within the fibril core. Phenylalanine and Tyrosine residues are indicated in magenta and salmon. Alanine substitutions are colored green. **(F)** Experimental workflow for monitoring fibril formation by ThT fluorescence over approximately 7 days. **(G)** Aggregation T_1/2_ values derived from the ThT kinetic measurements for Y275A, Y283A and F309A substitutions and the combined Y275A/Y283A/F309A mutant (Core_Triad_). Data are shown as an average across three technical replicates with error bars representing standard deviation. Core_F309A_ and Core_Triad_ both slowed fibril formation compared to WT. **(H)** Experimental workflow used to assess fibril stability after 14 days of incubation followed by ultracentrifugation and separation of supernatant and pellet fractions. **(I)** Concentration of hnRNPA2 remaining in the supernatant after ultracentrifugation for WT and the indicated core mutants. Increased soluble protein indicates reduced fibril stability. Data are shown as averages across three replicates with error bars representing standard deviation. **(J)** Representative negative-stain electron micrographs of endpoint samples from WT, Core_Y275A_, Core_Y283A_, Core_F309A_ and Core_Triad_ reactions. Core_Triad_ predominantly formed poorly organized or amorphous assemblies rather than the extensive fibrillar structures observed for WT and the single mutants.

### The aromatic triad is predicted to stabilize hnRNPA2 fibrillar state

We next used computational alanine scanning to define the energetic contributions of individual residues to this conserved amyloid fold [36, 37]. Three-, six- and nine-layer fibril assemblies were minimized and subjected to backrub sampling before systematic alanine substitution across the ordered core (Fig. 1C-D). Following minimization, the wild-type (WT) and mutant ensembles remained close to the experimental structure across all three assembly sizes, with RMSDs <0.4 Å for both the WT and alanine-substituted models (SI Appendix, Fig. 3A-C). This indicated that the energetic changes predominantly reflected local differences in packing rather than large-scale deformation of the fibril. To capture the changes in energetics in response to a mutation across the three-, six- and nine-layer size assemblies we interpreted the energetics using two different paradigms: 1) Estimating total differences in energy between WT and mutant assemblies (Fig. 1C; ΔREU_mut-wt_) and 2) Estimating the changes in energetic contribution of interfaces between WT and mutant assemblies (Fig. 1D and SI Appendix, Fig. 4A; ΔΔG_interface_). Across the complete scan, the magnitude of ΔREU_mut-wt_ destabilization upon alanine mutation increased with assembly size and varied strongly by residue class (SI Appendix, Fig. 4B-C). Glycine substitutions produced some of the largest changes in ΔREU_mut-wt_, particularly in the larger assemblies, whereas aromatic Phe/Tyr substitutions formed a distinct set of destabilizing positions (SI Appendix, Fig. 4B-C). Glycine-to-alanine substitutions produce large ΔREU_mut-wt_ changes due to steric clashes introduced upon mutation by the alanine methyl group, generating a van der Waals (vdW) penalty that correlates with ΔREU_mut-wt_ (SI Appendix, Fig. 4D, fa_rep). By contrast, mutation of aromatic residues to alanine leads to destabilization of ΔREU_mut-wt_ due to a loss of favorable vdW interactions (SI Appendix, Fig. 4E, Δfa_atr). Compared to ΔREU_mut-wt_, ΔΔG_interface_ showed stronger enrichment of aromatic residues among mutations that disrupted interactions between fibril layers, while the larger glycine to alanine mutation penalties were largely normalized (Fig. 1E-F and SI Appendix, Fig. 4F). These results distinguish glycines that are broadly sensitive to substitution from aromatic residues that make repeated, fibril-specific packing contacts across the amyloid assembly.

**Figure 3.**
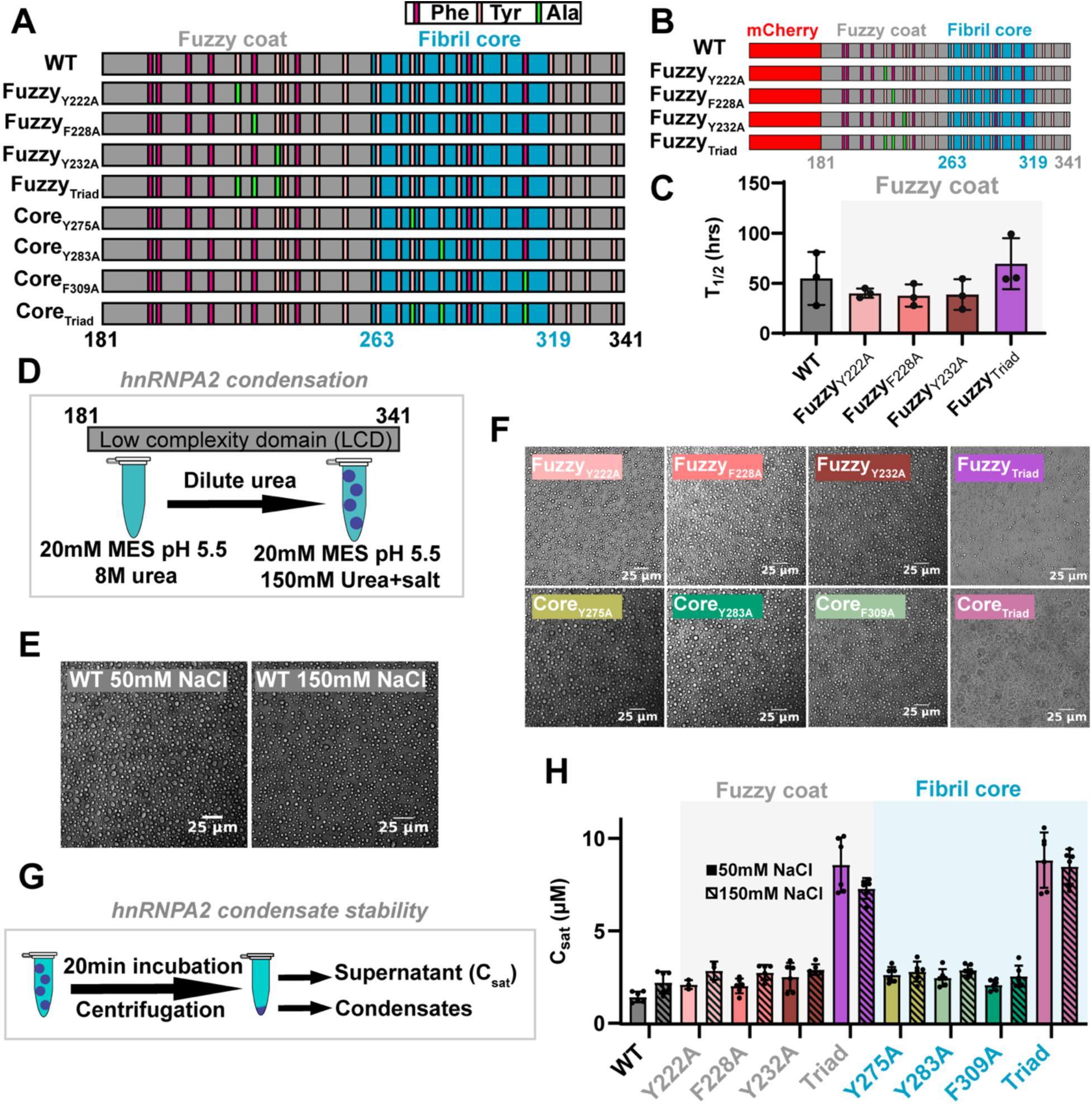
Alanine mutations alter phase separation similarly between the fuzzy coat and fibril core regions. **(A)** Schematic of hnRNPA2-LCD constructs containing aromatic-to-alanine substitutions either within the fuzzy coat (Fuzzy_Y222A_, Fuzzy_F228A_, Fuzzy_Y232A_ and Fuzzy_Triad_) or within the fibril core (Core_Y275A_, Core_Y283A_, Core_F309A_ and Core_Triad_). Phe, Tyr and introduced Ala residues are indicated. Cartoon is colored as in Fig. 2E. **(B)** Schematic of mCherry-hnRNPA2-LCD constructs containing aromatic-to-alanine substitutions in the fuzzy coat (Fuzzy_Y222A_, Fuzzy_F228A_, Fuzzy_Y232A_ and Fuzzy_Triad_). Phe, Tyr and introduced Ala residues are indicated. Cartoon is colored as in Fig. 2E. **(C)** Aggregation T_1/2_ values derived from the ThT kinetic measurements for mCherry-hnRNPA2-LCD WT and Fuzzy coat mutants (WT, Fuzzy_Y222A_, Fuzzy_F228A_, Fuzzy_Y232A_ and Fuzzy_Triad_). Data are shown as mean ± SD from three technical replicates. **(D)** Experimental scheme for inducing hnRNPA2-LCD phase separation by dilution from 8 M urea into low-urea MES buffer containing salt. **(E)** Representative light-microscopy images of WT hnRNPA2-LCD condensates formed at 50 or 150 mM NaCl. Scale bars, 25 μm. **(F)** Representative light-microscopy images of condensates formed by the indicated fuzzy-coat and fibril-core aromatic mutants. Scale bars, 25 μm. **(G)** Experimental workflow for determination of the saturation concentration (C_sat_). Following phase separation and a 20-min incubation, condensed and dilute phases were separated by centrifugation and protein concentration in the supernatant was measured. **(H)** C_sat_ values for WT and the indicated fuzzy-coat and fibril-core mutants at 50 and 150 mM NaCl. Data are shown as an average across two biological replicates each in triplicate (n=6) except Y222A (n=3) with errors represented as standard deviation.

**Figure 4.**
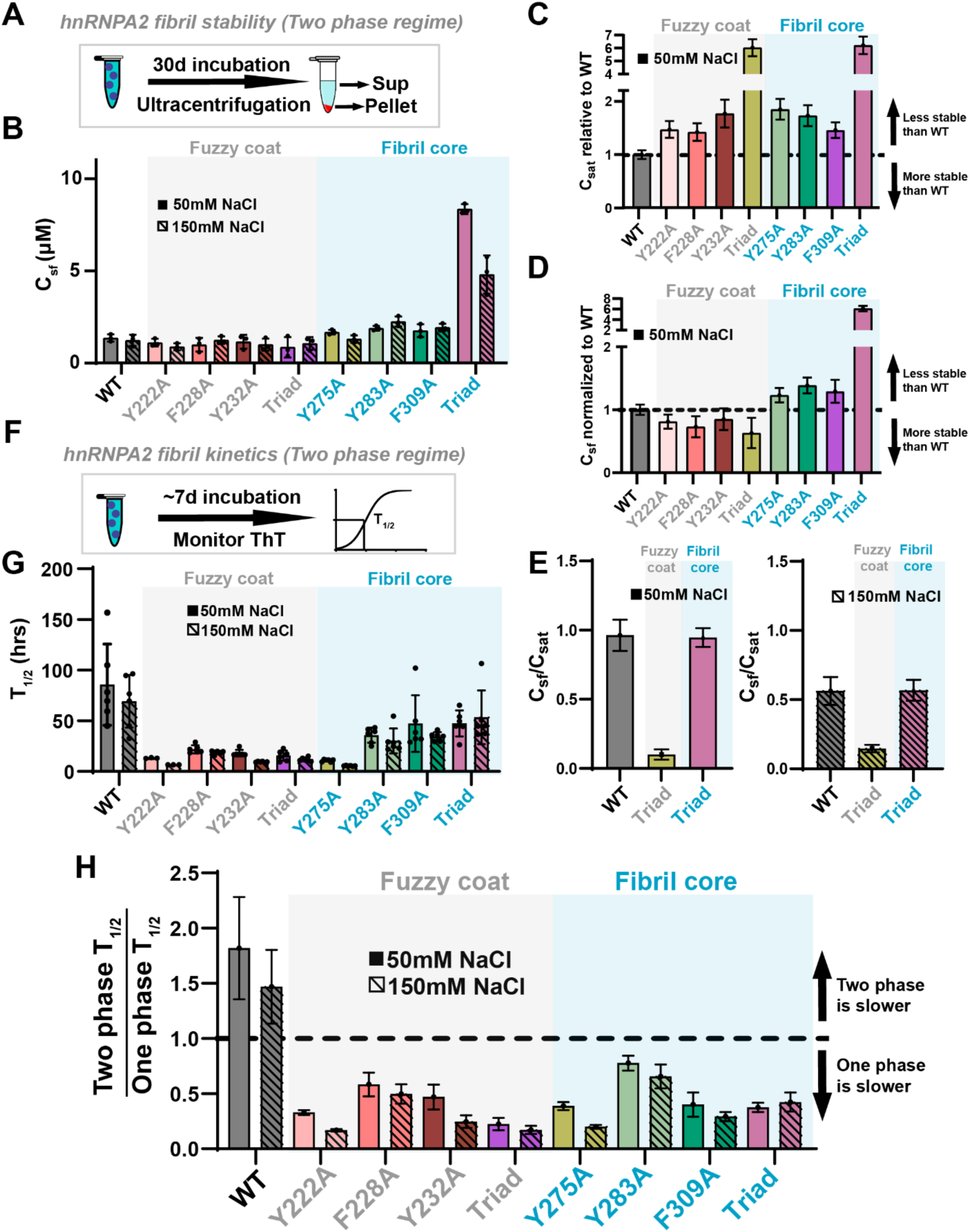
Aromatic substitutions differentially affect condensate stability, fibril stability and aggregation kinetics. **(A)** Experimental workflow used to determine fibril stability in the two-phase regime. Phase separation was initiated; samples were incubated for 30 days and soluble and fibrillar fractions were separated by ultracentrifugation. **(B)** Fibril solubility concentration (C_sf_) for WT and the indicated fuzzy-coat and fibril-core mutants at 50 and 150 mM NaCl. Data are shown as an average across three technical replicates with errors represented as standard deviation. **(C)** C_sat_ normalized to WT, for constructs incubated at 50 mM NaCl. Values above 1 indicate condensates are less stable, whereas values below 1 indicate condensates are more stable. Data are shown as an average with errors represented as SEM. **(D)** C_sf_ normalized to WT, for constructs incubated at 50 mM NaCl. Values above 1 indicate fibrils are less stable, whereas values below 1 indicate fibrils are more stable. Data are shown as an average with errors represented as SEM. **(E)** C_sf_/C_sat_ ratios for WT and Triad mutants at 50 mM and 150 mM NaCl. Data are shown as an average across three measurements with error bars representing SEM. **(F)** Experimental schematic for monitoring fibril formation of hnRNPA2 in the two-phase regime by ThT fluorescence over approximately 7 days. **(G)** Aggregation half-times (T_1/2_) for WT and fuzzy-coat and fibril-core mutants in the two-phase regime at 50 and 150 mM NaCl. Data are shown as averages across triplicates across 2 biological replicates (n=6) (except Y222A, n=3) with errors representing standard deviation. **(H)** Ratio of T_1/2_ measured in the two-phase versus one-phase regime for WT, fuzzy coat and fibril core mutants. Values below 1 indicate slower aggregation in the one-phase regime, whereas values above 1 indicate slower aggregation after phase separation. Data are shown as an average with errors represented as SEM.

The aromatic residues showed a particularly defined energetic pattern across assembly sizes. Mapping total and interface energetic changes for Phe/Tyr positions identified Y275, Y283, F309 and Y319 among the strongest aromatic hotspots, with the effects increasing from three- to six- to nine-layer assemblies (Fig. 1G-H and SI Appendix, Fig. 5A-B). Structural mapping of Y275, Y283 and F309 places these residues within the densely packed fibril core (Fig. 1I). Y319 on the other hand localizes to the C-terminus and forms favorable multilayer interactions with itself and forms a cross-layer hydrogen bond to Q267 (SI Appendix, Fig. 5C). The Y275, Y283 and F309 positions were particularly notable because they form a spatially connected packing network: F309 from one layer packs against Y275 and Y283 in the adjacent layer, directly linking the aromatic cluster to interlayer stabilization (Fig. 1J). Consistent with this structural arrangement, F309 showed a pronounced interface-energy loss upon alanine substitution, suggesting F309 forms interactions between layers while Y275 and Y283 primarily contributed to intralayer interactions in the aromatic triad (Fig. 1E, G, H). Analysis of the three aromatic substitutions individually and in combination reinforced the cooperative contribution of this network. Across three-, six- and nine-layer models, Y275A, Y283A and F309A each increased total and interface energies but the combined Y275A/Y283A/F309A triad produced a substantially larger energetic loss than any single substitution, with the difference becoming most pronounced in the six- and nine-layer assemblies (Fig. 1G, H and SI Appendix, Fig. 5A-B). The same trend was observed when ΔREU_mut-wt_ and ΔΔG_interface_ were compared with changes in solvent-accessible surface area: single substitutions occupied a relatively compact range, whereas the triad shifted toward both larger ΔSASA_mut-wt_ and larger total and interface energetic penalties, particularly for the larger assemblies (SI Appendix, Fig. 5D-E). Together, these analyses identify the Y275/Y283/F309 aromatic triad in the amyloid core as a cooperative packing hotspot that stabilizes the hnRNPA2 amyloid fold through repeated intra- and interlayer contacts and provides a focused set of residues for experimentally testing how fibril-specific interactions control hnRNPA2 aggregation.

**Figure 5.**
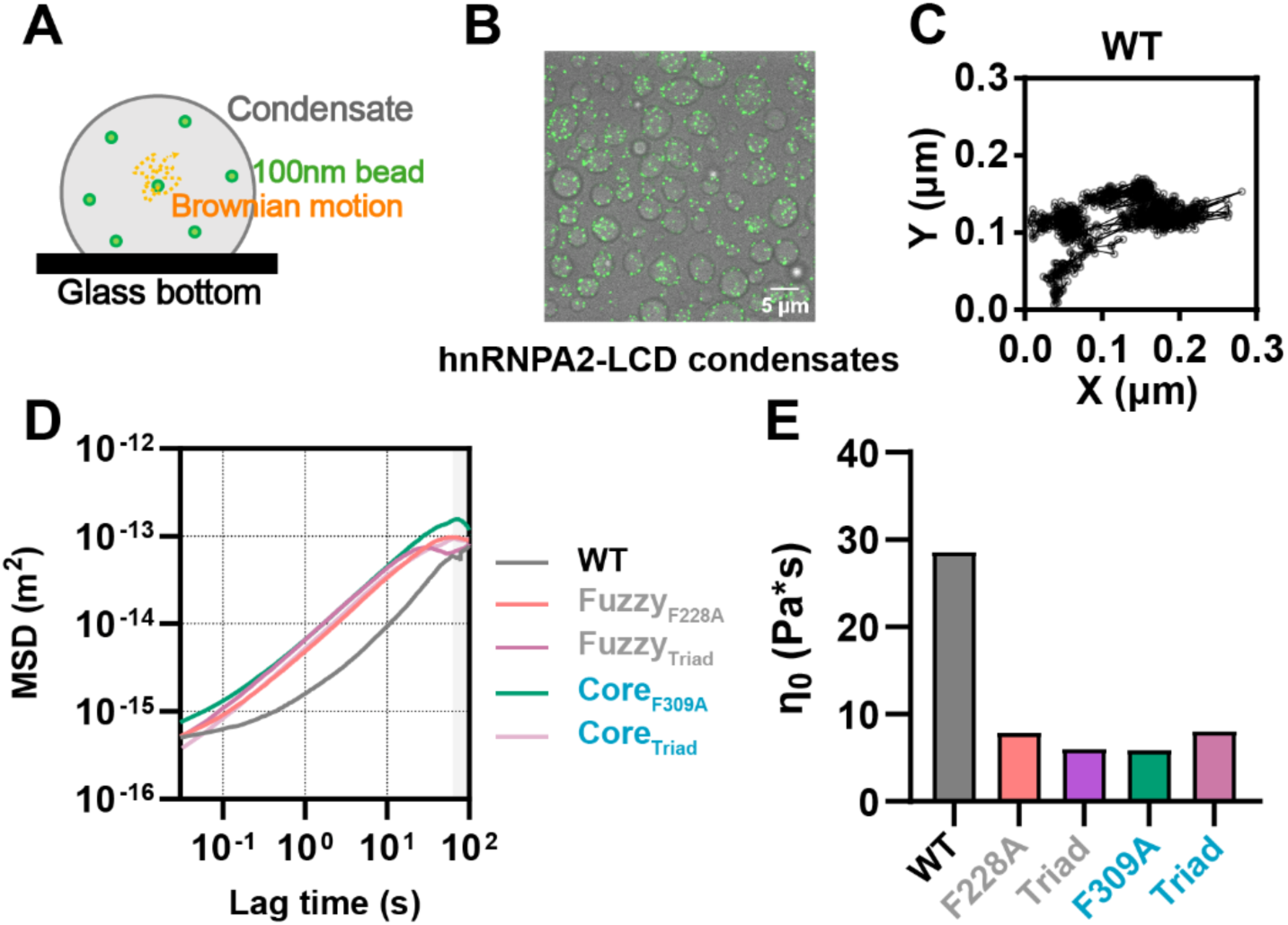
WT condensates are more viscous than fuzzy and fibril core mutants. **(A)** Schematic of video particle-tracking microrheology. Fluorescent 100-nm beads embedded within hnRNPA2-LCD condensates undergo thermally driven Brownian motion that reports on the local material properties of the condensate. **(B)** Representative fluorescence image of hnRNPA2-LCD condensates containing embedded tracer beads. Scale bar, 5 μm. **(C)** Representative two-dimensional trajectory of a tracer bead within a WT hnRNPA2-LCD condensate. **(D)** Ensemble-averaged mean-squared displacement (MSD) as a function of lag time for WT, Fuzzy_F228A_, Fuzzy_Triad_, Core_F309A_ and Core_Triad_ condensates. Reduced MSD in WT condensates indicates greater restriction of probe motion. Grey box represents region not used for further GSER analysis due to high error. **(E)** Zero-shear viscosity (η_0_) derived from particle-tracking microrheology measurements. WT condensates exhibit substantially higher viscosity than the aromatic mutant condensates.

### The Y275/Y283/F309 aromatic network controls hnRNPA2 amyloid formation and stability

To experimentally test the aromatic packing interactions identified by our structural and computational analyses, we first examined a series of peptides spanning different regions of the hnRNPA2 fibril core to determine the minimal interactions necessary for fibril formation. The longest peptide encompassed residues 273-311 and included all three residues of the Y275/Y283/F309 aromatic triad thereby spanning most of the central β- arch of the amyloid core that also includes the previously described amyloidogenic motif ^286^GNYNDF^291^ that carries disease-associated mutations (Fig. 2A and SI Appendix, Fig. 6A; herein Full turn). We additionally generated a series of truncations that systematically removed portions of this structural element and altered which aromatic residues were retained (Fig. 2A and SI Appendix, Fig. 6A). Orthogonally, we used sequence-based amyloid propensity predictions using ZipperDB[38] which identified several contiguous high-propensity segments within the hnRNPA2 fibril core, including regions encompassing the Y275/Y283 aromatic residues and the amyloidogenic ^286^GNYNDF^291^ hexapeptide (SI Appendix, Fig. 6B)[9]. Fibril formation was monitored by Thioflavin-T (ThT) fluorescence [39]. The full turn peptide encoding residues 273-311 formed ThT positive flourescence rapidly with a T_1/2_ of 21.32±0.26 hrs (SD) (Fig. 2B, SI Appendix, Fig. 6C and 7A), while partial turn peptide encoding 281-311, which retains Y283 and F309 but lacks Y275, formed amyloid even faster with a T_1/2_ of 6.63±0.202 hrs (SD) (Fig. 2B, SI Appendix, Fig. 6C and 7B). By contrast, peptide fragments lacking either Y283 or F309 showed variability between replicates and little or no increase in ThT fluorescence (Fig. 2B, SI Appendix, Fig. 6C and SI Appendix, Fig. 7C; half turn, N-term half and C-term half) with the predicted longer amyloidogenic segment showing slow kinetics with a low final fluorescence amplitude (Fig. 2B, SI Appendix, Fig. 6C and SI Appendix, Fig. 7C; ZipperDB). Thus, among the fragments tested, the partial turn encoding residues 281-311 represents the minimal sequence capable of efficient amyloid formation, indicating that the combination of Y283 and F309 is sufficient to support assembly whereas Y275 is dispensable in this peptide context. Importantly, our data also support that the core element in the hnRNPA2 LCD that controls aggregation must contain both clustering of aromatic residues and the aggregation-prone hexapeptide region[9, 38].

Using our peptide system, we next tested the effect of alanine substitutions at each position within the aromatic triad. We used the full turn peptide (residues 273-311) due to it spanning the entire aromatic triad (Fig 2C). We incubated each peptide with ThT and monitored fluorescence (Fig. 2D, SI Appendix, Fig. 6D and 8A-E). Similar to previous findings, Y275A was comparable to WT fibril formation with a modest increase in T_1/2_ (31.94±2.56 hrs (SD) and 17.92±2.94 hrs (SD)), respectively while alanine substitutions at Y283, (96.11±36.25 hrs (SD)), F309 (116.28±40.22 hrs(SD)) and the Triad (no fit) (SI Appendix, Fig. 8A-E) had the greatest increase in T_1/2_ supporting that the clustering of aromatics within the peptide, not the amyloid motif, drives aggregation.

We next tested the contribution of these aromatic residues in the context of the hnRNPA2 LCD using the same _mCherry-hnRNPA2-LCD_ construct used for structural analysis. Individual Y275A, Y283A and F309A substitutions were introduced into the fibril core (Fig. 2C, herein Core_Y275A_, Core_Y283A_ and Core_F309A_), together with a combined Y275A/Y283A/F309A mutant (Fig. 2E, herein Core_Triad_). Following removal of denaturing buffer, fibril formation was monitored by ThT fluorescence over seven days (Fig. 2F). No evidence of phase separation was seen during incubation with low turbidity for all constructs. Increases in turbidity at later timepoints were correlated with increases in ThT fluorescence and formation of fibrils which can contribute to turbidity (SI Appendix, Fig 6E-F). Aggregation half-times showed that the mutations that retained detectable aggregation assembled more slowly than WT (39.40 ± 8.10 hrs (SD)) except Core_Y275A_ (27.80±3.90 hrs (SD)) (Fig. 2G and SI Appendix, Fig. 9A-C). The mutants with the strongest kinetic defects were Core_F309A_ (118.07±26.79 hrs (SD)) and Core_Triad_ (126.57±5.52 hrs (SD)) whose ThT fluorescence signals did not reach saturation during the experiment (SI Appendix, Fig. 9D). These results indicate that the contributions of the aromatic residues are not equivalent: Y275 is largely dispensable for fibril formation, whereas F309 is particularly important and simultaneous disruption of the aromatic triad strongly impairs assembly.

We next asked whether these same interactions contribute to the stability of the assembled fibril. Following a 14 day incubation, samples were subjected to ultracentrifugation and the amount of soluble protein remaining in the supernatant was quantified by densitometry (Fig. 2H). WT and Core_Y275A_ remained predominantly in the insoluble fraction, while Core_Y283A_ produced a slight increase in soluble protein. By contrast, Core_F309A_ showed a substantial increase in the soluble fraction and Core_Triad_ exhibited the greatest loss of fibril stability, with the majority of protein remaining in the supernatant (Fig. 2I). Negative-stain electron microscopy of endpoint samples was consistent with these measurements. WT, Core_Y275A_, Core_Y283A_ and Core_F309A_ samples contained recognizable fibrillar assemblies, whereas Core_Triad_ was dominated by short, poorly organized or amorphous structures rather than the extensive fibrils observed for WT (Fig. 2J). Together, these data demonstrate that the central aromatic packing network identified in the hnRNPA2 fibril structure is important for amyloid assembly and stability. Among the three residues, F309 is the only one that contributes to interlayer contacts, and combined disruption of Y275, Y283, and F309 strongly impairs formation of the native hnRNPA2 amyloid architecture.

### Aromatic contributions to hnRNPA2 phase separation are largely independent of their position within the LCD

Having established that aromatic residues within the fibril core strongly influence the rate of formation and stability of the hnRNPA2 amyloid, we next asked how these same residues contribute to hnRNPA2 liquid-liquid phase separation. To examine condensate formation independently of the mCherry solubility tag, we used the isolated hnRNPA2 LCD and generated matched aromatic-to-alanine substitutions either within the fuzzy coat outside the ordered fibril core (Y222A, F228A and Y232A; Fuzzy mutants) or within the fibril core (Y275A, Y283A and F309A; Core mutants) (Fig. 3A). We selected these fuzzy coat residues based on previous reports that their serine substitutions do not affect hnRNPA2 hydrogel partitioning[32]. We additionally generated combined triple mutants at each region, termed Fuzzy_Triad_ and Core_Triad_ (Fig. 3A).

To establish whether the fuzzy-coat substitutions alter aggregation in the absence of condensates, we also introduced these mutations into the mCherry-hnRNPA2-LCD construct used for the one-phase experiments (Fig. 3B). All three single mutants and Fuzzy_Triad_ retained ThT-positive aggregation, with aggregation half-times broadly comparable to WT (Fig. 3C and SI Appendix, Fig. 10A-F). These substitutions therefore did not produce the pronounced kinetic defects observed for Core_F309A_ and Core_Triad_ (Fig. 2G), providing a baseline for evaluating their effects under phase-separated conditions.

Phase separation was initiated using an established dilution protocol in which hnRNPA2 LCD was transferred from 8 M urea into low urea buffer containing either 50 or 150 mM _NaCl_ [40] (Fig. 3D). WT hnRNPA2 rapidly formed spherical droplets within seconds at both 50 and 150 mM NaCl (Fig. 3E). Likewise, all individual Fuzzy and Core aromatic mutants, as well as both triad mutants, retained the ability to form visible condensates (Fig. 3F and SI Appendix, Fig. 11A). Thus, neither individual aromatic substitutions nor simultaneous disruption of three aromatic residues was sufficient to abolish phase separation under these conditions.

We next quantified the contribution of these residues to condensate stability by measuring the saturation concentration (C_sat_). Following phase separation, samples were incubated for 20 min and centrifuged to separate the condensed and dilute phases and the concentration of protein remaining in the supernatant was measured as C_sat_ (Fig. 3G). Within this timeframe and up to one hour, there is little evidence of fibril formation with no exponential growth in ThT signal and no visible fibrils (SI Appendix, Figs. 11B and 17A-B). Increasing the salt concentration from 50 to 150 mM generally increased C_sat_, consistent with reduced condensate stability at higher ionic strength (Fig. 3H). Individual aromatic-to-alanine substitutions produced comparatively slight increases in C_sat_ and this effect was similar whether the substituted residue was located within the fuzzy coat or the fibril core. By contrast, simultaneously replacing three aromatic residues produced a substantially larger increase in C_sat_. Both FuzzyTriad and Core_Triad_ exhibited the strongest defects in phase separation, with similarly elevated C_sat_ values at both salt concentrations (Fig. 3H). These results indicate that aromatic residues contribute collectively to hnRNPA2 condensate stability but that this contribution is not strongly dependent on whether the aromatic residues reside within the ordered amyloid core or the surrounding fuzzy coat. Thus, whereas the fibril experiments identify specific core aromatics, particularly F309 and the Y275/Y283/F309 network, as critical determinants of amyloid assembly, condensate formation appears to depend more broadly on the overall aromatic valence of the LCD. This distinction allowed us to next determine whether disrupting the fibril-specific aromatic network alters amyloid formation when hnRNPA2 is first assembled within a two-phase condensate regime.

### The core aromatic triad coordinately destabilizes condensate and fibril states of hnRNPA2

We next asked how aromatic substitutions within the fuzzy coat or fibril core alter the transition from phase-separated condensates to fibrils. Phase separation was initiated as described above and samples were incubated for 30 days before ultracentrifugation to quantify the concentration of protein remaining soluble at equilibrium with the fibrillar state (C_sf_) (Fig. 4A). At 50 mM NaCl, mutations within the fuzzy coat produced relatively small decreases in C_sf_ compared with WT, indicating that these residues have only a minor effect on stability of the mature fibril state (Fig. 4B). Individual substitutions within the fibril core only increased around 1.4 −1.8 fold in C_sf_ compared to control fuzzy mutants, whereas Core_Triad_ increased around 9-fold in C_sf_ compared to Fuzzy_Triad_, consistent with fibril destabilization (Fig. 4B). This pattern persisted at 150 mM NaCl, although the destabilizing effect of Core_Triad_ was reduced at the higher ionic strength (around 4.5 fold increase, Fig. 4B). Notably, Core_F309A_ behaved similarly to the other single core mutants, in contrast to its stronger destabilizing effect in the one-phase experiments. Thus, the combined Y275/Y283/F309 aromatic network remains a major determinant of fibril stability when amyloid formation proceeds through a phase-separated intermediate.

To compare the effects of aromatic substitutions on condensate and fibril stability, we normalized C_sat_ and C_sf_ to their respective WT values (Fig. 4C-D and SI Appendix, Fig. 12A-B). At 50 mM NaCl, substitutions in either region increased C_sat_, with Fuzzy_Triad_ and Core_Triad_ producing the largest increases. Their effects on C_sf_ differed: fuzzy-coat substitutions decreased C_sf_, whereas individual core substitutions produced modest increases and Core_Triad_ produced the largest increase. A similar overall pattern was observed at 150 mM NaCl. Comparing C_sf/_C_sat_ further distinguished the two triad mutants (Fig. 4E and SI Appendix, Fig. 12C-D). Fuzzy_Triad_ exhibited a substantially lower ratio than WT at both salt concentrations, reflecting increased C_sat_ together with reduced C_sf_. By contrast, Core_Triad_ retained a ratio close to WT at each salt concentration, consistent with approximately proportional increases in condensate and fibril solubility. These results indicate that the effects of aromatic substitutions on fibril stability depend strongly on their location within the ordered core, whereas their effects on condensate stability are less dependent on the region examined.

### Aromatic substitutions accelerate fibril formation under two-phase conditions

We next determined how aromatic substitutions influence the kinetics of condensate-to-fibril conversion. Following induction of phase separation, fibril formation was monitored by ThT fluorescence for around 7 days and aggregation half-times (T_1/2_) were determined (Fig. 4F-G). The isolated WT hnRNPA2 LCD aggregated relatively slowly (85.60±40.16 hrs (SD)), whereas all aromatic alanine mutants displayed shorter T_1/2_ values under the two-phase conditions (Fig. 4G and SI Appendix, Figs. 13A-C, 14A-G, 15A-H and 16A-B). The strongest acceleration was observed for fuzzy-coat mutants and Core_Y275A_ (Fuzzy_Y222A_ 13.18±0.34 hrs (SD), Fuzzy_F228A_ 21.95±3.93 hrs (SD), Fuzzy_Y232A_ 18.12±3.20 _hrs_ (SD), Core_Y275A_ 10.77±1.16 hrs (SD), Fuzzy_Triad_ 15.58±5.13 hrs (SD)). By comparison, Core_Y283A_ (35.81±7.54 hrs (SD)), Core_F309A_ (47.22±27.85 hrs(SD)) and Core_Triad_ (47.45±12.86 hrs(SD)) formed fibrils more slowly than their fuzzy-coat counterparts, consistent with a specific requirement for the core aromatic network during fibril assembly (Fig. 4G). Increasing the ionic strength from 50 to 150 mM generally accelerated fibril formation, although WT and Core_Triad_ were comparatively insensitive to changes in ionic strength. Direct imaging supported these kinetic measurements where the droplets formed by the fuzzy-coat mutants and Core_Y275A_ rapidly developed fluorescent amyloid structures, whereas Core_Y283A_, Core_F309A_ and particularly Core_Triad_ retained droplet-like morphologies for longer periods (SI Appendix, Figs. 17A-B and Movie S1-S18).

We next compared aggregation half-times across the one-phase and two-phase assays for WT and all single and triple aromatic mutants (Fig. 4H). The fuzzy-coat mutants, whose one-phase aggregation half-times were broadly comparable to WT (Fig. 3C and SI Appendix, Fig. 10), displayed pronounced acceleration relative to WT under two-phase conditions (Fig. 4G). Across the two assay formats, all aromatic mutants had two-phase/one-phase half-time ratios below one, whereas WT had ratios above one. Together, these observations support a contribution from the condensate environment to the accelerated aggregation of the fuzzy-coat mutants. This pattern prompted us to examine whether the mutations alter the material properties and molecular organization of hnRNPA2 condensates.

#### Increased condensate viscoelasticity is associated with slower fibril conversion

WT hnRNPA2 displayed strikingly different fibrilization kinetics depending on whether aggregation occurred in a one-phase or two-phase regime. In the one-phase regime, WT hnRNPA2-LCD formed fibrils with relatively rapid kinetics, whereas under the two-phase regime, WT assembled more slowly than either fuzzy-coat or fibril-core aromatic mutants (Figs. 2G and 4G). This suggested that properties emerging from the dense phase, in addition to intrinsic fibril-forming propensity, could regulate the condensate-to-fibril transition [27, 41, 42]. To test this possibility, we used video particle-tracking microrheology (VPT) to measure the material properties of hnRNPA2-LCD condensates. We introduced inert fluorescent 100-nm beads to condensates and the thermally driven bead motions were tracked over time (Fig. 5A-C). Condensates were stabilized on the treated glass surface to minimize bulk movement and trajectories from beads outside or near the condensate boundary were excluded. The resulting bead trajectories provided a local readout of the material properties of the hnRNPA2-LCD network.

We first compared the mean-squared displacement (MSD) of beads within WT condensates and representative fuzzy-coat and fibril-core mutants. One hour after phase separation, beads in Fuzzy_F228A_, Fuzzy_Triad_, Core_F309A_ and Core_Triad_ condensates showed substantially greater motion than beads in WT condensates across the measured lag times (Fig. 5D). The reduced motion in WT condensates could be explained by the elastic constraint uniquely present in WT in the measured time scale. Consistent with this idea, WT showed subdiffusive motion due to elastic constraint at shorter lag times (slope α < 1), whereas bead motion in the mutant condensates more closely approached Brownian diffusion (slope α = 1), indicating that the WT dense phase forms a more mechanically restrictive network.

We next transformed the MSD measurements into frequency-dependent viscoelastic moduli using the generalized Stokes-Einstein relation assuming a simple Maxwell fluid behavior (see Methods). WT and mutant condensates displayed viscoelastic behavior characterized by frequency-dependent storage (G’) and loss (G’’) moduli (SI Appendix, Fig. 18A-E). At low frequencies, the viscous component dominated, whereas the elastic component dominated at higher frequencies. Importantly, the slopes of G’ and G” are ∼1 and ∼2, which is a characteristic of G’ and G” of Maxwell fluids at terminally viscous regions. The inverse of the frequency at the crossover between G’ and G’’ provided a characteristic relaxation time for the condensate network. WT condensates exhibited a substantially longer relaxation time than either Fuzzy_F228A_ or Core_F309A_, while the relaxation times of Fuzzy_Triad_ and Core_Triad_ were shorter than the accessible timescale (SI Appendix, Fig. 18F). These measurements indicate that at the 100-nm scale, the network of WT condensates takes longer to rearrange and behaves like an elastic solid in time scales where the network of aromatic mutants freely rearranges and behaves like a viscous fluid.

The difference in condensate dynamics was also reflected in the zero-shear viscosity, η_0_, which is a measurement independent of timescale. WT condensates exhibited a substantially higher η_o_ than the aromatic mutants, approximately 28.6 Pa·s compared with ∼6-8 Pa·s for Fuzzy_F228A_, Fuzzy_Triad_, Core_F309A_ and Core_Triad_ (Fig. 5E). Thus, removal of aromatic interactions from either the fuzzy coat or fibril core markedly reduced the zero-viscosity of the condensate. Importantly, this effect was independent of the location of the substituted residues, consistent with our C_sat_ measurements showing that aromatic residues contribute broadly to condensate organization rather than through a fibril-core-specific interaction network. Together, these results reveal a strong correspondence between condensate material properties and fibrilization kinetics. WT hnRNPA2 forms a comparatively viscous condensate in which molecular diffusion is reduced, coinciding with its unusually slow conversion to fibrils in the two-phase regime. Aromatic substitutions reduce condensate viscosity and increase predicted molecular mobility, while simultaneously accelerating fibril formation. These observations support a model in which viscoelasticity of the WT hnRNPA2 condensate acts as a kinetic barrier to amyloid formation by restricting the molecular rearrangements and dilute and dense phase exchange required for productive fibril assembly.

## Discussion

Extensive work has examined how condensate stability influences the transition from phase separation to amyloid formation, yet considerably less is known about how the material properties of condensates and the structural energetics of the amyloid state act together to control this transition. Here, using the structurally reproducible hnRNPA2 low-complexity domain (LCD) system, we combined cryo-EM, computational energetic profiling, peptide design and LCD mutational analysis with measurements of aggregation propensity, phase behavior and condensate microrheology to define how these two states are coupled. We identify a central aromatic packing network involving Y275, Y283 and F309 that makes a disproportionate contribution to the mature fibril fold, with F309 emerging as a particularly important interaction hub in the one-phase condition that we have shown forms a reproducible amyloid fold. The pronounced destabilizing effect of Core_F309A_ in the one-phase assay was less evident under two-phase conditions, where the individual core substitutions produced comparatively modest changes in fibril stability. Core_F309A_’s differing behavior across phase regimes may reflect distinct experimental conditions, with condensates potentially favoring an alternative amyloid conformation. Core_Triad_ nevertheless strongly destabilized the fibrillar state in both experimental regimes, supporting an important collective contribution from the Y275/Y283/F309 network. Differences in construct and solution conditions may contribute to the altered sensitivity to individual substitutions; whether these differences also involve changes in fibril architecture remains unresolved. At the same time, our phase-separation experiments reveal that aggregation propensity is not determined solely by the stability of the amyloid state: WT hnRNPA2 aggregates rapidly outside the condensate regime but is comparatively inhibited in fibril formation within condensates, whereas under two-phase conditions, all tested aromatic mutants exhibited shorter aggregation half-times than WT, with the strongest acceleration observed for the fuzzy-coat mutants and Core_Y275A_. Microrheology provides a likely physical explanation for this behavior, showing that WT condensates form a more constrained, viscoelastic network with higher viscosity and longer relaxation times, while mutant condensates examined had lower viscosities and exhibited more liquid-like behavior over the measured timescales. Together, these findings support a model in which hnRNPA2 amyloid formation is governed by the interplay between two separable but coupled energetic landscapes: condensate material properties determined by aromatic residues throughout the sequence kinetically regulate access to the fibrillar state, while a precisely organized aromatic network in the fibrillar core determines the stability of the mature amyloid structure. Thus, aromatic interactions can either restrain aggregation by establishing a viscoelastic condensate environment or promote aggregation by stabilizing the amyloid core, depending on their structural context.

### Amyloid-core and fuzzy-coat aromatics contribute similarly to phase separation

A striking relationship emerged when the same aromatic interactions were examined in the phase-separated state. Aromatic substitutions within the fibril core and matched substitutions within the surrounding fuzzy coat produced similar increases in saturation concentration and the Core_Triad_ and Fuzzy_Triad_ mutants had comparably strong effects on condensate stability (Fig. 3). These results are consistent with sticker-and-spacer models of phase separation in which aromatic residues function as distributed interaction sites whose collective valence and patterning promote condensation[21]. In this context, whether a particular aromatic residue ultimately becomes buried within the mature amyloid core appears to be less important than its contribution to the overall interaction network of the disordered LCD.

Our findings, however, argue against a simple model in which the molecular interactions stabilizing the condensed and amyloid states are equivalent. Core_Triad_ disrupts three residues that become buried and structurally integrated within the fibril and consequently has a strong effect on fibril stability. By contrast, Fuzzy_Triad_ alters aromatic residues outside the ordered amyloid core and has relatively little effect on the stability of the mature fibril. Nevertheless, both perturbations similarly weaken phase separation. Thus, the condensate and fibrillar states rely on overlapping chemical features but organize those interactions in fundamentally different ways. Aromatic valence appears to be an important determinant of condensation, whereas the precise spatial arrangement of specific aromatic residues becomes critical in the amyloid state. This distinction is also evident when condensate and fibril stability are considered together. Individual mutations within the amyloid core can preferentially destabilize the fibrillar state relative to the condensate, whereas fuzzy-coat mutations preferentially alter phase separation. Remarkably, simultaneous disruption of Y275, Y283 and F309 destabilizes both states sufficiently that the relative balance between condensate and fibril stability approaches that of WT. We therefore interpret the Core_Triad_ phenotype not as evidence that the two states use identical interactions but rather that a single sequence perturbation can affect two partially decoupled energetic minima to similar extents.

### Condensate viscoelasticity establishes a kinetic barrier to amyloid formation

The kinetic behavior of hnRNPA2 revealed an additional level of regulation that could not be explained by fibril thermodynamics alone. WT hnRNPA2 readily formed fibrils from a homogeneous solution, yet fibril formation was markedly slower when the protein first assembled into condensates. In contrast, aromatic substitutions within either the fuzzy coat or fibril core generally accelerated fibril formation in the two-phase regime. Thus, simply increasing the local protein concentration through phase separation is not sufficient to accelerate amyloid formation. Instead, the physical properties of the condensed phase appear to determine whether condensation promotes or suppresses the transition to the amyloid state. Microrheology provided a likely mechanistic explanation for this behavior. WT hnRNPA2 condensates were substantially more viscous and exhibited longer relaxation times than condensates formed by the fuzzy-coat and core aromatic mutants. Mahendran, Gui and Bremer et al. proposed a model in which the highly viscoelastic condensates create a kinetic barrier by restricting protein efflux into the dilute phase which limits fibril elongation at the condensate periphery [29]. Our hnRNPA2 data are consistent with a model in which the WT condensate interaction network limits protein exchange with the dilute phase and thereby slows fibril elongation. Reduced viscosity in the aromatic mutants could relieve this constraint, although protein efflux and individual microscopic steps of fibril assembly were not directly measured. Importantly, condensate fluidity alone does not determine the rate of amyloid formation. Core_Triad_ and Fuzzy_Triad_ produce condensates with similarly altered material properties, yet Core_Triad_ converts to fibrils substantially more slowly. This difference is readily explained by the fibril kinetics: Fuzzy_Triad_ lowers the viscoelastic kinetic barrier without substantially compromising the interactions that kinetically drive amyloid formation, whereas Core_Triad_ simultaneously lowers condensate viscosity, but loss of the aromatic triad slows fibril formation. Thus, condensate material properties determine how efficiently hnRNPA2 molecules encounter and explore aggregation-competent configurations, while fibril-specific interactions determine whether those configurations can mature into a stable amyloid structure. Amyloid kinetics therefore emerge from the combination of condensate-scale molecular mobility and fibril-scale structural compatibility.

### Reconciling protective and aggregation-promoting models of biomolecular condensates

This framework may help reconcile apparently contradictory observations concerning the relationship between biomolecular condensation and amyloid formation. In hnRNPA1 and related systems, strengthening condensate interactions or introducing additional condensate-stabilizing components has been reported to suppress fibril formation[26, 29]. Such findings support a model in which the condensed phase can act as an inhibitory state. Conversely, condensates and stress-granule-like assemblies formed by proteins such as TDP-43 have been proposed to serve as environments that facilitate pathological aggregation[43]. Our results suggest that these models need not be mutually exclusive. Whether a condensate promotes or suppresses amyloid formation may depend strongly on its material state. A highly viscoelastic condensate can concentrate protein and initiate fibril nucleation at the interface while simultaneously restricting the molecular diffusion and structural rearrangements required for fibril elongation[29]. In this model, a more fluid condensate may retain the concentration advantage of phase separation while permitting faster protein exchange between the dense and dilute phases, potentially supporting fibril elongation following nucleation at the interface [23, 24, 29]. The important variable may therefore not simply be whether a protein undergoes phase separation but the mobility, relaxation dynamics and interaction lifetimes present within the dense phase. This model also suggests that the material properties of disease-associated condensates may help determine their propensity to mature into pathological aggregates. Condensates that remain highly viscoelastic could kinetically trap proteins in a reversible dense phase, whereas perturbations that increase molecular mobility could promote structural conversion even without dramatically altering the equilibrium concentration of the protein. Testing this idea across different aggregation-prone proteins will be important for determining whether condensate rheology represents a general regulator of pathological phase transitions.

### Y275 may function as a kinetic gate to amyloid formation

Y275 is particularly interesting because its effect on aggregation differs from that of Y283 and F309. Although Y275 contributes to the mature fibril architecture, Y275A substantially accelerates aggregation in both one-phase and two-phase conditions. Similarly, the shorter peptide lacking the N-terminal Y275-containing segment of the fibril turn aggregates more rapidly than the corresponding longer peptide. These observations suggest that Y275 may influence the kinetic accessibility of the amyloid fold rather than functioning simply as a favorable stabilizing interaction in the final structure. One potential explanation involves the local environment surrounding D276. Y275 lies immediately adjacent to D276, which occupies an interlayer environment within the fibril, raising the possibility that removal of the Y275 side chain alters the geometry or energetic frustration associated with D276. A related phenomenon has been described for D290, where _changes_ in the local interlayer environment can influence hnRNPA2 aggregation behavior[44]. Y275A could therefore lower a local kinetic barrier by relieving unfavorable packing or electrostatic constraints around D276, allowing the fibril architecture to form more readily. This interpretation remains a structural hypothesis rather than a demonstrated mechanism. Direct energetic decomposition of the Y275/D276 environment or structural analysis of Y275A fibrils will be necessary to determine whether altered D276 interactions account for the accelerated kinetics. Nevertheless, the unusual behavior of Y275 emphasizes that residues incorporated into a mature amyloid structure can have different effects on the kinetics and thermodynamics of amyloid assembly.

### A multiscale model for the hnRNPA2 condensate-to-fibril transition

Together, our findings support a multiscale model in which the hnRNPA2 condensate-to-fibril transition is controlled by two interacting but separable energetic landscapes. At the mesoscale, distributed aromatic interactions throughout the LCD determine condensate stability, viscosity and relaxation dynamics. These properties define the kinetic environment in which amyloid nucleation occurs. At the molecular scale, structurally specific interactions within the fibril core determine whether aggregation-competent configurations can mature into the stable hnRNPA2 amyloid architecture. WT hnRNPA2 combines a highly viscoelastic condensate with a strongly stabilized fibril fold. The dense phase therefore concentrates hnRNPA2 but simultaneously imposes a kinetic barrier to structural conversion. Removing aromatic interactions from the fuzzy coat weakens this viscoelastic network and accelerates molecular rearrangement without substantially altering the stability of the resulting amyloid state, thereby strongly accelerating condensate-to-fibril conversion. Mutations within the amyloid core likewise make the condensate more fluid but also alter the ability of the sequence to fold into an amyloid conformation. The Core_Triad_ represents the most extreme example because it disrupts the viscoelastic protection of the condensate while simultaneously removing a central packing network required for efficient formation and stability of the mature amyloid.

This model provides a framework for understanding why condensation can either inhibit or promote amyloid formation depending on sequence and context. The condensate is not simply an intermediate of elevated protein concentration, nor is the fibril an inevitable thermodynamic endpoint of phase separation. Instead, the transition reflects competition between distributed interactions that define condensate material properties and structurally encoded interactions that define the amyloid fold. The interplay between these two scales determines both the energetic preference for each state and the kinetic accessibility of the pathway connecting them.

## Data availability

The generated cryo-EM datasets are available on the Electron Microscopy Data Bank repository under accession numbers EMD-78776 and EMD-78777. Additionally, the cryo-EM model is available on the Protein Data Bank under PDB IDs 38FN and 38FP. Protein energetic calculations, aggregation kinetics and thermodynamics data, condensate data, microscopy images and VPT microrheology data are available as source data. Raw data associated with these data are available on Zenodo under accession number 22818452.

## Code availability

Software used for determining the hnRNPA2-LCD structure includes RELION v4.0(https://github.com/3dem/relion-documents), CTFFIND v4.1 [45], COOT v0.8.9 (https://www2.mrc-lmb.cam.ac.uk/personal/pemsley/coot/). All energetic calculations on the hnRNPA2-LCD fibril structure were performed using an adapted flex_ddg protocol available on https://git.biohpc.swmed.edu/s184069/flex_ddg_ala_scn_runner using Rosetta v3.12 https://www.rosettacommons.org/. VPT microrheology and microscopy images were analyzed by Fiji v2.16.0 (https://imagej.net/software/fiji/downloads) and MATLAB. Graphs were generated using GraphPad Prism v11.0.2.

## Supporting information

see SI Appendix, Movie legends

see SI Appendix, Movie legends

see SI Appendix, Movie legends

see SI Appendix, Movie legends

see SI Appendix, Movie legends

see SI Appendix, Movie legends

see SI Appendix, Movie legends

see SI Appendix, Movie legends

see SI Appendix, Movie legends

see SI Appendix, Movie legends

see SI Appendix, Movie legends

see SI Appendix, Movie legends

see SI Appendix, Movie legends

see SI Appendix, Movie legends

see SI Appendix, Movie legends

see SI Appendix, Movie legends

see SI Appendix, Movie legends

see SI Appendix, Movie legends

## Acknowledgements

L.A.J. was supported by an Effie Marie Cain Scholarship in Medical Research and a grant from the NIH-NIA (RF1AG076459). M.K.R. was supported by the Howard Hughes Medical Institute and grants from the Welch Foundation (I-1544) and NIH (R35GM141736). Transmission electron microscopy was performed at the UT Southwestern Electron Microscopy Core Facility, supported by the National Institutes of Health (1S10OD021685-01A1 and 1S10OD020103-01). Cryo-EM data were acquired at the Cryo-Electron Microscopy Facility (CEMF) and the Structural Biology Laboratory (SBL) at UT Southwestern, which are supported by a grant from the Cancer Prevention & Research Institute of Texas (RP220582). Computational resources were provided by the BioHPC cluster supported by the Lyda Hill Department of Bioinformatics at UT Southwestern. Special thanks to Dr. Kunach for guidance on cryo-EM structure determination, the McKnight lab for the mCherry-hnRNPA2 plasmid and the Joachimiak lab for helpful discussions and feedback on the manuscript.

## Competing interests statement

The authors declare no competing interests.

## Author Contributions

D.J.K. and L.A.J. initiated and conceived of the project. D.J.K. determined the structure of the hnRNPA2-LCD. D.J.K. conducted all aggregation related experiments and conducted condensate stability and imaging experiments. J.H.H. conducted all VPT microrheology experiments. D.J.K. and L.A.J. wrote the manuscript and all authors contributed to the revisions.

## Methods

### Plasmid generation

The WT mCherry-hnRNPA2 construct was provided by the McKnight lab at University of Texas, Southwestern Medical Center. Point mutants were generated by either site-directed mutagenesis or synthesized gene blocks from Twist Biosciences or Integrated DNA Technologies (IDT). The DNA sequence encoding the mCherry was removed by PCR and ends were re-ligated to generate untagged hnRNPA2 constructs.

### Purification of mCherry-hnRNPA2 and untagged hnRNPA2 constructs

mCherry-hnRNPA2 was purified similarly as discussed previously[33]. Protein was overexpressed in One Shot BL21 (DE3) pLysS cells (Thermofisher scientific) with 0.5mM IPTG overnight at 16C. Pellets were suspended with 50 mM Tris pH 8.0, 500 mM NaCl, 2M Urea, 2mM TCEP, 1X Roche protease inhibitor (Millipore Sigma) and stored at −80C. Pellets were sonicated and spun at 14,000x g for 30 min to remove insoluble material. The resulting supernatants were incubated with HisPur NiNTA resin (Thermofisher) for 30 min and added to a gravity column (Biorad). NiNTA bound mCherry-hnRNPA2 was washed with 25mM Tris pH 8.0, 500 mM NaCl, 2M Urea, 1mM PMSF, 20 mM Imidazole and 2mM TCEP until A280 reached baseline. mCherry-hnRNPA2 was then eluted with 25mM Tris pH 8.0, 500 mM NaCl, 2M Urea, 1mM PMSF, 200 mM Imidazole and 2mM TCEP. mCherry-hnRNPA2 was then concentrated using an Amicon 10 kDa MWCO Ultra Centrifugal Filter (Millipore sigma) to either 10mg/mL for fibril kinetics and stability assays or 60-80mg/mL for hydrogel formation. Aliquots were finally snap frozen and stored at 80°C.

Untagged hnRNPA2-LCD was purified as described previously[40]. hnRNPA2 was overexpressed in One Shot BL21 (DE3) pLysS until OD reached 0.8-1.0 and induced with 1mM IPTG at 25C overnight. Harvested pellets were resuspended with 25 mM Tris-HCl (pH 8), 500 mM NaCl, 2mM TCEP and 1X Roche protease inhibitor and stored at −80C. Pellets were sonicated and spun at 14,000x g for 30 min to separate insoluble material. The hnRNPA2 containing pellet was resuspended with 25 mM Tris-HCl, 500 mM NaCl, 10 mM imidazole, 1mM DTT, 8M Urea rocking at RT for 1hr. Extracted hnRNPA2-LCD was separated from cell debris by centrifugation (30,000x g for 30 min). Supernatants were added to HisPur NiNTA resin (Thermofisher) and incubated for 20 min. The resin was then washed with ∼10CV of 25 mM Tris-HCl, 500 mM NaCl, 15 mM imidazole, 1mM DTT, 8M Urea. hnRNPA2-LCD was eluted with 25 mM Tris-HCl, 500 mM NaCl, 300 mM imidazole, 1mM DTT and 8M urea and concentrated to 1mL final volume using an Amicon 10 kDa MWCO Ultra Centrifugal Filter (Millipore sigma). For TEV cleavage, concentrated hnRNPA2 was diluted 1:10 with TEV cleavage buffer (25mM Tris pH8, 50 mM NaCl, 1mM DTT). Final urea concentration 800 mM. hnRNPA2 was cleaved overnight with TEV protease at RT. Crashed out hnRNPA2 was resuspended with 20 mM Tris pH 7.5 and 8M urea until resuspended and concentrated to 300µM. Aliquots were flash frozen and stored at −80C

### Formation of mCherry-hnRNPA2 hydrogels and fibrils

mCherry-hnRNPA2 constructs were thawed and filtered with a 0.22µm spin filter (Thermo fisher) before dialysis. Constructs were dialyzed at RT in 20 mM Tris pH 7.5 200 mM NaCl 20 mM BME .5mM EDTA .1mM PMSF overnight. Crashed out protein was removed by centrifugation (14,000x g for 15min). Soluble protein was incubated at RT (stability and kinetics) or incubated at 4C for hydrogel formation.

### Cryo-EM data acquisition

mCherry-hnRNPA2-LCD hydrogels were diluted to ∼0.5mg/mL and 2.5uL was applied to glow-discharged Quantifoil Cu R1.2/1.3, 300 mesh carbon grids. Samples were blotted using Vitrobot Mark IV (FEI) at 4C, 100% humidity. Data was collected on a 300kV Titan Krios using a Gatan K3 Summit detector in CDS mode. The total electron dose was 60 e-/Å^2 with a pixel size of 0.738Å. To eliminate inelastically scattered electrons, a GIF quantum energy filter (Gatan) with a slit width of 20 eV was employed.

### Helical reconstruction

Collected movies were motion corrected and dose weighted using the motion correction implementation provided by RELION 4.0[46]. Motion corrected micrographs were then used to estimate their contrast transfer function (CTF) using CTFFIND-4.1[45].

Particle picking was done automatically using Topaz[47]. The resulting particles were extracted with a 450Å (612px) box size with a 14.25Å interbox distance and then downscaled by a factor of 6. These particles were used for the first round of reference free 2D classification using a mask diameter of 440Å and a tube diameter of 100Å. The highest resolution 2D classes were chosen. The respective particles were then re-extracted with a 280Å (388px) box size with no rescaling and another round of reference free 2D classification was run. The highest resolution 2D classes were then used for 3D classification.

The first round of 3D classification was run using the density map from previous work[31] that was low pass filtered to 10Å and resampled to match pixel size. Particles for final round of refinement were polished and CTF refined. The final resolution estimate was 1.99Å at the gold standard 0.143 FSC cutoff[48].

To determine the structure of the polymorph, we did an additional round of 3D classification with the particles used for initial structure determination which generated a new 3D class not shown in the previous refinements. The generated reference map and particles from the class were refined, polished and CTF corrected. The final resolution estimate was 2.12 Å at the 0.143 FSC cutoff.

Both maps were built in coot using PDB 6wqk as an input model[31]. Newly resolved residues within the polymorph were built into the map manually. The models were refined using Phenix 1.20 [49]

### In silico alanine scan

Varying numbers of layers of fibril protofilaments were prepared for use in silico estimation of assembly energies as described previously[37]. To summarize, hnRNPA2 fibril structure was retrieved from the Protein Data Bank (PDB). Using Pymol (version 3.1), fibril PDB structures were created with numbers of layers varying between three and nine. Changes in assembly energy were calculated using a method adapted from Flex ddg[36] for amyloid fibrils[37]. For both the interface and the *ΔREU^assembly^* calculations, all chains were mutated. For the ΔΔ*G*^interface^ calculations on 3,6 and 9-mer assemblies presented, the chains of either the center 1, 2 or 3 layers for each respective structure were used to define the interface. Using the fa_talaris2014 score function, the structure then underwent minimization. After minimization, the residues within 8 angstroms of the mutation site underwent backrub sampling and with were then repacked and minimized. The Rosetta InterfaceDdgMover was used as in the Flex ddG protocol to allow an analysis of the ΔΔ*G*^interface^ by defining a fibril interface, giving the lowest energy bound and unbound states for both a wildtype and mutant fibril. For *ΔREU^assembly^*calculations, the bound wild-type and bound mutant structures reported by the interface ddG mover were used for estimating the change in assembly energy due to an alanine substitution. This is repeated for 35 independent replicates and repeated for every residue in the structure to generate a set of *ΔREU^assembly^* values which were then averaged. To calculate ΔΔ*G*^interface^ for a residue, the Flex ddG protocol was used as described previously[36]. Briefly, the Δ*G* of binding was calculated by subtracting the bound state energy from the unbound state energy for both a wildtype and mutant fibril and the Δ*G*_wt_ was subtracted from the Δ*G*_mut_ to yield the ΔΔ*G*^interface^ for that alanine mutant. This was repeated for every residue to generate a set of ΔΔ*G*^interface^ values which were averaged over 35 structures.

### RMSD analysis

Backbone RMSD analysis of the output structure was carried with an in-house script that extracts structures using a Rosetta database file generated by the Flex ddG protocol. The C-α RMSDs were then calculated using the align command in PyMOL. RMSD distributions were visualized using GraphPad version 11.0.2.

### Peptide synthesis and disaggregation

Peptides were synthesized by Genscript with N-terminal acetylation and C-terminal amidation modifications at >95% purity. Peptides were disaggregated in 250uL Trifluoroacetic acid (TFA) (Thermo Fisher) and incubated overnight in a fume hood to allow for full evaporation. Peptides were resuspended with 200uL H_2_O, flash frozen and lyophilized. The resulting powder was resuspended with 20 mM Tris pH 7.5, 200 mM NaCl, 20 mM BME, .5mM EDTA, .1mM PMSF and 8M urea and snap frozen in 1.5mM-2mM stock. Peptides were stored in −80°C until use.

### Thioflavin-T aggregation assay of hnRNPA2 peptides

Peptide concentrations were adjusted to 1066.67µM with 20 mM Tris pH 7.5, 200 mM NaCl, 20 mM BME, .5mM EDTA, .1mM PMSF and 8M urea. Aggregation was initiated by diluting the peptide into 20 mM Tris pH 7.5 200 mM NaCl 20 mM BME .5mM EDTA .1mM PMSF to a final urea concentration of 150 mM (20µM final protein concentration). Thioflavin-T was added at a final concentration of 20µM. Technical triplicates of 15uL were aliquoted into a 384 low profile microplate (Aurora) and sealed with Crystal clear plate seals (Hampton research). The measurements were recorded at an excitation of 446 nm and emission of 486 nm at 15min intervals in a Tecan Spark plate reader at quiescent conditions. The data points were plotted as averages with standard deviation. The T_1/2_ was estimated for each replicate using a non-linear sigmoid fit in GraphPad Prism v11.2. Each technical replicate was fitted and reported as an average of three replicates with standard deviation.

### Thioflavin-T kinetics and fibril stability of mCherry-hnRNPA2-LCD fibrils and mutants

Thioflavin-T (ThT) was added to mCherry-hnRNPA2 constructs (20µM ThT, final protein concentration 20µM) and added in triplicate to 384 low profile microplates (Aurora). Plates were sealed with Crystal clear plate seals (Hampton research) and spun at 1000g for 5min to remove bubbles. Plates were inserted into a Tecan Spark plate reader. The measurements were recorded at an excitation of 446 nm and emission of 486 nm at 15min intervals. The data points were plotted as averages with standard deviation. The T_1/2_ was estimated for each replicate using a non-linear sigmoid fit in GraphPad Prism v11.2. Each technical replicate was fitted and reported as an average of three replicates with standard deviation.

For thermodynamic measurements, 20µM mCherry-hnRNPA2-LCD was incubated at 25°C in a thermoshaker (Eppendorf) for 14 days. Samples were transferred to thick-walled ultracentrifuge tubes (Beckman Coulter) and spun at 186,000g for 30 min. The supernatant was removed and the pellet was resuspended with 8M urea. Equal volumes of sample were run on a SDS-PAGE gel and stained with SimplyBlue stain (Thermo Fisher). Band intensity was quantified using Fiji. % in supernatant was quantified by dividing the amount of protein left in supernatant by the amount of protein in pellet+supernatant. The percentage was multiplied by the starting protein concentration to get final protein concentration in supernatant.

### Fibril preparation for negative stain transmission electron microscopy

4uL of fibrils were added to glow discharged 400mesh formvar grids and adhered for 2.5 minutes. Grids were washed for 30secs with water and stained with 4uL uranyl acetate for 2min. Excess stain was blotted off and grids were imaged using a Tecnai G2 Spirit or a JEOL 1400 Plus electron microscope operating at 120 kV provided by the UT Southwestern Medical Center Electron Microscopy Core facility.

### Untagged hnRNPA2-LCD phase separation

Untagged hnRNPA2-LCD constructs were desalted into 20 mM MES pH 5.5 and 8M urea using a Zeba 7k MWCO desalting column. Constructs containing aggregates were filtered with a 0.02µm filter (Wyatt technology). Constructs were then concentrated to 1.5mM-2mM in a 10K MWCO spin filter. Constructs were diluted to 1066.67µM with 20 mM MES pH 5.5, 8M urea. Phase separation was initiated by diluting the urea into 20 mM MES pH 5.5 with either 50 mM or 150 mM NaCl buffer (final urea concentration 150 mM, 20µM final protein concentration).

### Saturation concentration determination of hnRNPA2-LCD constructs

hnRNPA2-LCD phase separation was initiated and incubated for 20 min at RT (final protein concentration 20µM). The resulting dilute and dense phases were separated by centrifugation (14,000x g, 20 min). The dilute phase was measured using a DeNovix Nanodrop by measuring absorbance at 280nm. Triplicates were plotted as averages with standard deviation (SD).

### Microscopy imaging of hnRNPA2 condensates

CellVis 384 glass bottom plates were passivated with 100mg/mL BSA as described previously[50] before droplet imaging. Phase separation was initiated similarly as above but 20µM of ThT was added. Phase separated constructs were added to passivated plate and imaged with a 20X water immersion optic in an ImageXpress confocal HT.ai system. Images were normalized and wavelengths merged using Fiji.

### hnRNPA2 two-phase regime aggregation kinetics

Phase separated hnRNPA2 was added to a BSA passivated CellVis 384 glass bottom plate in triplicate and ThT fluorescence was monitored in a Tecan Spark plate reader. The measurements were recorded at an excitation of 446 nm and emission of 486 nm at 10 min intervals. The data points were plotted as averages with standard deviation. The T_1/2_ was estimated for each replicate using a non-linear sigmoid fit in GraphPad Prism v11.2. Each technical replicate was fitted and reported as an average of replicates with standard deviation.

### hnRNPA2 two-phase regime fibril stability

Phase separated hnRNPA2 was incubated at 25C for 23 days and then shaken at 600rpm on a thermomixer (Eppendorf) for 7 days afterward. Samples were transferred to thick-walled ultracentrifuge tubes (Beckman Coulter) and spun at 186,000g for 30 min. Soluble protein concentration was measured based on 280nm absorbance in a Denovix nanodrop.

### Sample preparation for particle tracking microrheology

Samples were prepared in 384-well glass bottom plates (Brooks Life Science Systems: MGB101-1-2-LK-L). The plates were cleaned with 5% Hellmanex III (Hëlma Analytics), etched with 1 M KOH and siliconized with Sigmacote (Sigma-Aldrich). Prior to sample addition, individual wells were blocked with 1 mg/mL Bovine Serum Albumin (BSA), excess BSA was removed and further washed with 20 mM MES pH 5.5 buffer. hnRNPA2-LCD samples were prepared using 20 μM protein in buffer containing a final concentration of 20 mM MES pH5.5, 50 mM NaCl, 150 mM Urea and 0.00004% solids 100 nm-diameter FluoSpheres^TM^ Carboxylate-Modified Microspheres (Invitrogen: F8803). Wells were sealed and samples were incubated inside the temperature-controlled microscope chamber at 23.5-28 °C for at least 1 hour before imaging.

### Microscopy imaging for particle tracking microrheology

Microscopy was performed on a spinning disk confocal (SDC) setup with a Leica DMI6000 B microscope base, a Yokogawa CSU-X1 scanner unit, a 405/488/561/647 nm Laser Quad Band Set filter cube (Chroma) and a plan apo 100 × 1.40 NA oil immersion objective. Images were captured with a Hamamatsu ImagEMX2 EM-CCD camera at 15-32 ms/frame using stream acquisition function on Metamorph (Biovision). 10000 frames (150-320 s) were acquired for each acquisition and multiple acquisitions were made per sample for reproducibility.

### Data analysis pipeline for particle tracking microrheology

Particle tracking and calculation of mean-squared displacement (MSD) were performed using MATLAB codes by Daniel Blair & Eric Dufresne (https://site.physics.georgetown.edu/matlab/code.html). Average MSD was calculated from <50,000 individual trajectories and smoothed using a moving average with span <10% of total number of frames. Viscous (G”) and elastic (G’) moduli as a function of frequency (ω) were calculated from the averaged MSD using Generalized Stokes-Einstein Relation (GSER) as described by Mason TG[51]

### Calculation of zero-shear viscosity, and terminal relaxation time

Viscosity (η) as a function of frequency (ω) was calculated from viscous modulus (G”) using η = G”/ ω. Low frequency region of the viscosity plot was then used to calculate zero-shear viscosity (η_0_). To calculate the terminal relaxation time, the inverse of the frequency at the crossover (G” = G’) was used.

## Author Contributions

D.J.K. and L.A.J. initiated and conceived of the project. D.J.K. determined the structure of the hnRNPA2-LCD. D.J.K. conducted all aggregation related experiments and conducted condensate stability and imaging experiments. J.H.H. conducted all microrheology experiments. D.J.K. and L.A.J. wrote the manuscript and all authors contributed to the revisions.

## Competing Interest Statement

All authors report no competing interests

---

**SI Appendix, Table 1:** Data collection, refinement, and validation statistics for hnRNPA2-LCD fibril structure.

| *Data collection and processing* | *hnRNPA2-LCD (PDB 38FN, EMD-78776)* |
|---|---|
| Magnification | ×138,000 |
| Voltage (kV) | 300 |
| Electron exposure (e<sup>−</sup> Å<sup>−2</sup>) | 60 |
| Defocus range (μm) | 0.9–2.2 |
| Pixel size (Å) | 0.738 |
| Symmetry imposed | C1; Helical |
| Initial particle images (no.) | 432,384 |
| Final particle images (no.) | 95,470 |
| Map resolution (Å) | 1.99 |
| FSC threshold | 0.143 |
| Map resolution range (Å) | 3.54–1.99 |
| *Refinement* |  |
| Initial model used | 6wqk |
| Model resolution (Å) | 2.05 |
| FSC threshold | 0.5 |
| Map sharpening *B* factor (Å<sup>2</sup>) | -27.7618 |
| *Model composition* |  |
| Non-hydrogen atoms | 2075 |
| Protein residues | 285 |
| *B factors (Å<sup>2</sup>)* |  |
| Protein | 29.48 |
| *R.m.s. deviations* |  |
| Bond lengths (Å) | 0.005 |
| Bond angles (°) | 1.031 |
| *Validation* |  |
| MolProbity score | 0.5 |
| Clashscore | 0 |
| Poor rotamers (%) | 0 |
| *Ramachandran plot* |  |
| Favored (%) | 98.18 |
| Allowed (%) | 1.82 |
| Disallowed (%) | 0 |

**SI Appendix, Table 2:** Data collection, refinement, and validation statistics for alternate N and C-termini confirmation of hnRNPA2-LCD fibril structure.

| *Data collection and processing* | *hnRNPA2-LCD alternate N and C termini conformation (PDB 38FP, EMD-78777)* |
|---|---|
| Voltage (kV) | 300 |
| Electron exposure (e<sup>−</sup> Å<sup>−2</sup>) | 60 |
| Defocus range (μm) | 0.9–2.2 |
| Pixel size (Å) | 0.738 |
| Symmetry imposed | C1; Helical |
| Initial particle images (no.) | 432,384 |
| Final particle images (no.) | 17,412 |
| Map resolution (Å) | 2.12 |
| FSC threshold | 0.143 |
| Map resolution range (Å) | 4.4043–2.12 |
| *Refinement* |  |
| Initial model used | PDB:6wqk |
| Model resolution (Å) | 1.92 |
| FSC threshold | 0.5 |
| Map sharpening *B* factor (Å<sup>2</sup>) | -26.0096 |
| *Model composition* |  |
| Non-hydrogen atoms | 2075 |
| Protein residues | 285 |
| *B factors (Å<sup>2</sup>)* |  |
| Protein | 33 |
| *R.m.s. deviations* |  |
| Bond lengths (Å) | 0.003 |
| Bond angles (°) | 0.97 |
| *Validation* |  |
| MolProbity score | 1.21 |
| Clashscore | 3.97 |
| Poor rotamers (%) | 0 |
| *Ramachandran plot* |  |
| Favored (%) | 97.87 |
| Allowed (%) | 2.13 |
| Disallowed (%) | 0 |

**SI Appendix, Figure 1.**
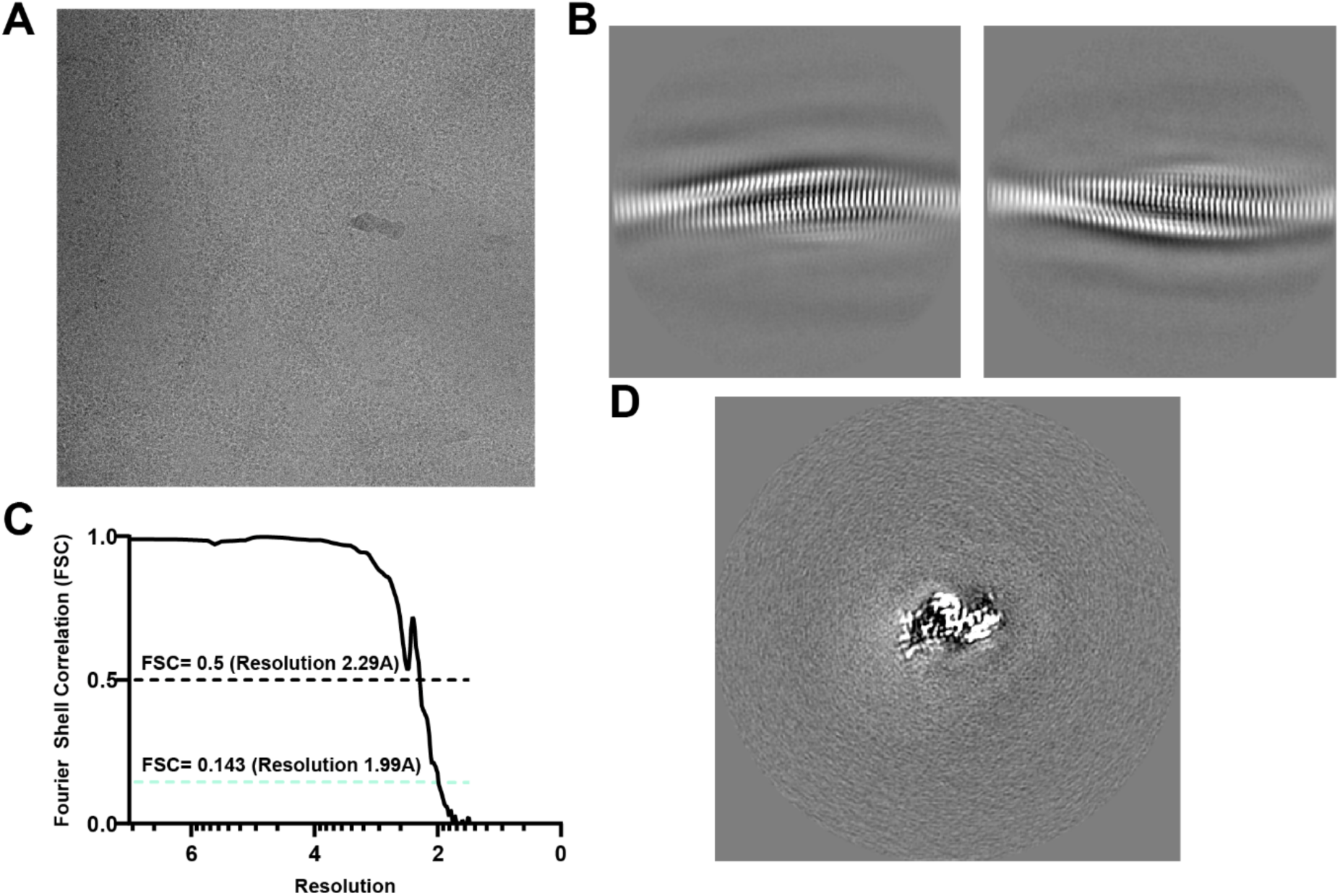
Cryo-EM reconstruction of the hnRNPA2 LCD fibril. **(A)** Representative cryo-EM micrograph of hnRNPA2 LCD fibrils used for structural determination. **(B)** Representative two-dimensional class averages showing the repeating cross-β architecture of the fibrils. **(C)** Fourier shell correlation (FSC) curve for the final reconstruction. The FSC = 0.143 criterion corresponds to an overall resolution of 1.99 Å; the FSC = 0.5 criterion is also indicated. **(D)** Representative view of the final cryo-EM reconstruction illustrating the ordered fibril density.

**SI Appendix, Figure 2.**
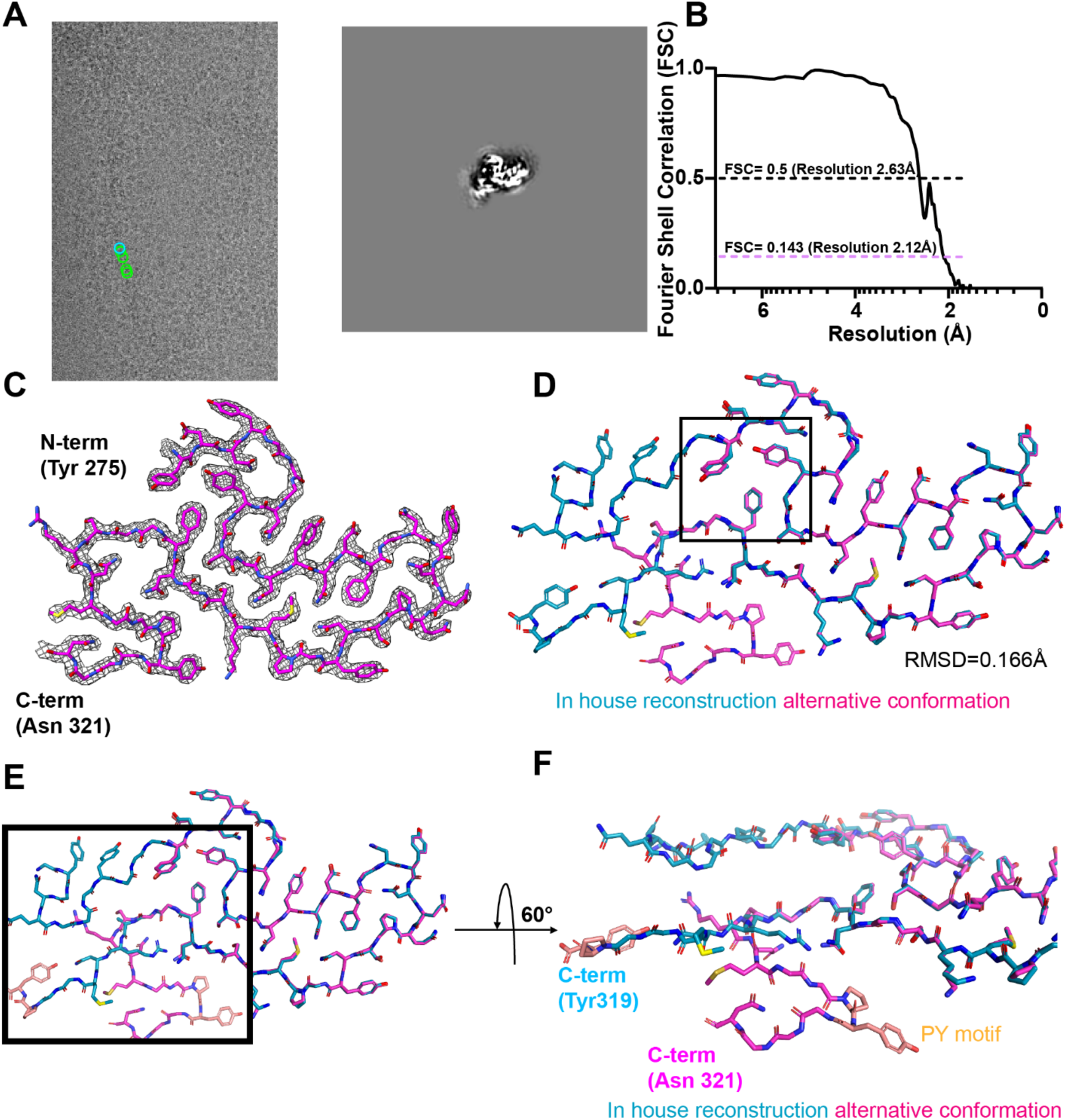
An alternate hnRNPA2 fibril conformation retains the central amyloid architecture. **(A)** Representative cryo-EM image and reconstructed projection of the alternate hnRNPA2 fibril population. Green circles and blue circles represent particles from the reproduced hnRNPA2 fibril structure and alternative structure respectively. **(B)** FSC curve for the alternate reconstruction, yielding an overall resolution of 2.12 Å at the FSC = 0.143 criterion. FSC = 0.5 criterion is also shown. **(C)** Cryo-EM density and atomic model for the alternate fibril conformation, with the ordered region extending from Y275 toward the C-terminal portion of the amyloid core. **(D)** Structural alignment of the alternate conformation with the primary hnRNPA2 reconstruction. The fibril architectures are highly similar, with an RMSD of 0.166 Å when aligning the core residues (275-309). **(E)** Comparison of the C-terminal portions of the primary and alternate fibril structures. **(F)** Rotated view highlighting differences in the C-terminal trajectory of the two structures and the positioning of residues near the PY-NLS-containing region. The central fibril fold remains conserved despite variation at the C-terminal edge of the ordered core.

**SI Appendix, Figure 3.**
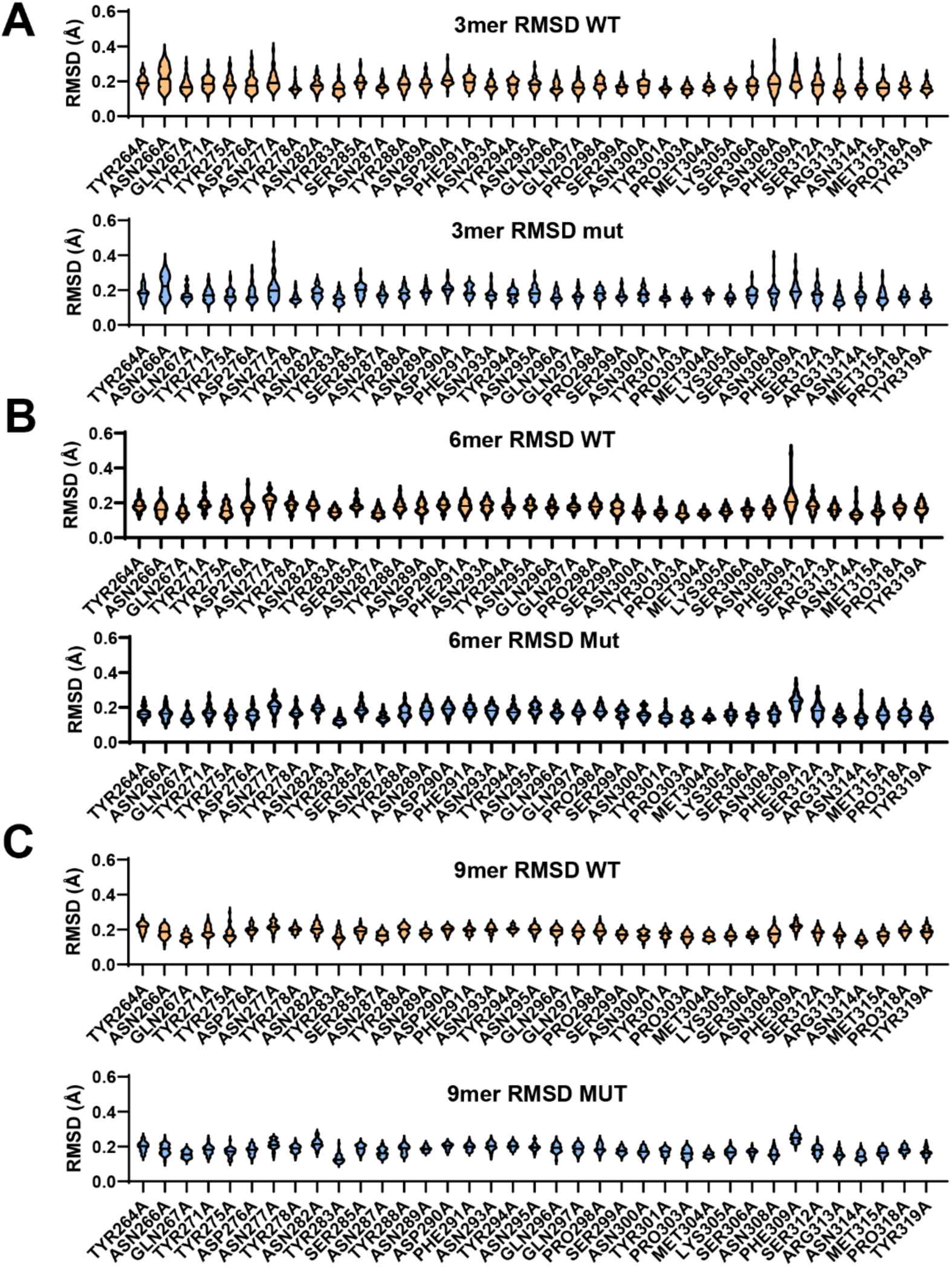
Rosetta sampling preserves the experimental hnRNPA2 fibril architecture. Distributions of backbone RMSD relative to the starting cryo-EM structure for WT and alanine-mutant structures generated during Rosetta minimization and backrub sampling. **(A)** Three-layer, **(B)** six-layer and **(C)** nine-layer fibril assemblies. Upper plots in each panel show RMSD distributions for the corresponding WT structures and lower plots show the alanine-mutant structures. Low RMSD values across the alanine scan indicate that the calculated energetic effects are not driven by large-scale disruption of the experimental fibril architecture. Data are shown as violin plots across 35 replicates.

**SI Appendix, Figure 4.**
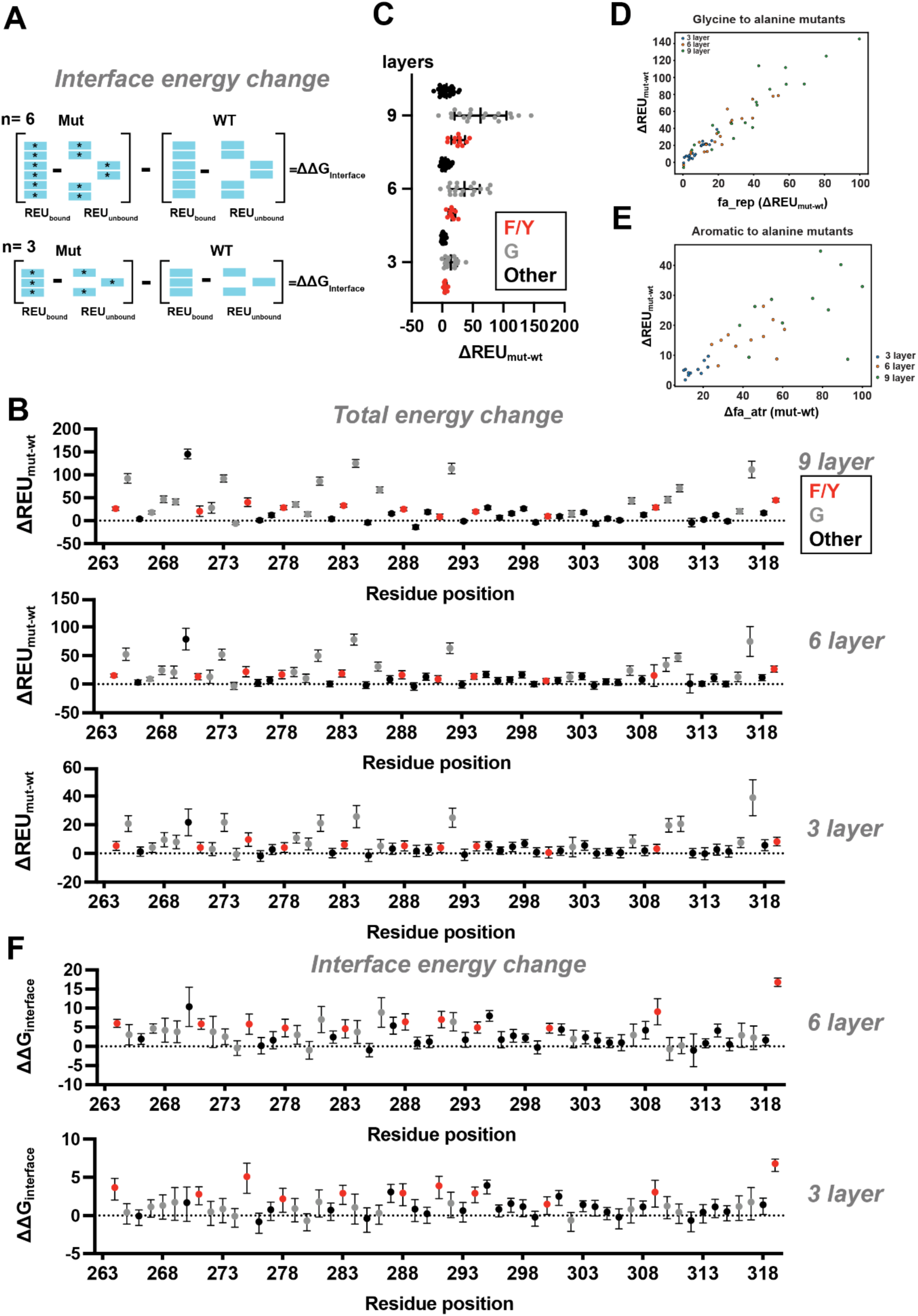
Energetic decomposition of the hnRNPA2 alanine scan identifies distinct contributions from glycine and aromatic residues. (A) Schematic illustrating calculation of the fibril interface energy. Energies of the bound and separated states are calculated independently for mutant and WT structures and their difference yields ΔΔGinterface. (B) Residue-by-residue total energetic changes (ΔREU_mut−WT_) following alanine substitution in 9-, 6- and 3-layer hnRNPA2 fibril assemblies. Aromatic residues (Phe/Tyr), glycines and other residues are colored red, grey and black. Points represent averages across 35 replicate calculations with standard deviation. (C) Distribution of total ΔREU values grouped by residue class and assembly size. Aromatic residues are shown in red, glycines in gray and other residues in black. Individual points are shown. Data is reported as an average with standard deviation. (D) Relationship between the fa_rep contribution and total energetic change for glycine-to-alanine substitutions in 3-, 6- and 9-layer assemblies in blue, orange and green. The strong relationship indicates that the large raw energetic penalties associated with Gly to Ala substitutions are dominated by steric repulsion introduced by the alanine methyl group. (E) Relationship between the change in the attractive van der Waals term (fa_atr) and total energetic change for aromatic-to-alanine substitutions in 3-, 6- and 9-layer assemblies in blue, orange and green. Loss of favorable fa_atr interactions contributes strongly to the energetic cost of removing Phe/Tyr side chains. (F) Residue-by-residue ΔΔG_interface_ values for the 6- and 3-layer fibril assemblies. Aromatic residues (Phe/Tyr), glycines and other residues are colored red, grey and black. Points represent averages across 35 replicate calculations with standard deviation.

**SI Appendix, Figure 5.**
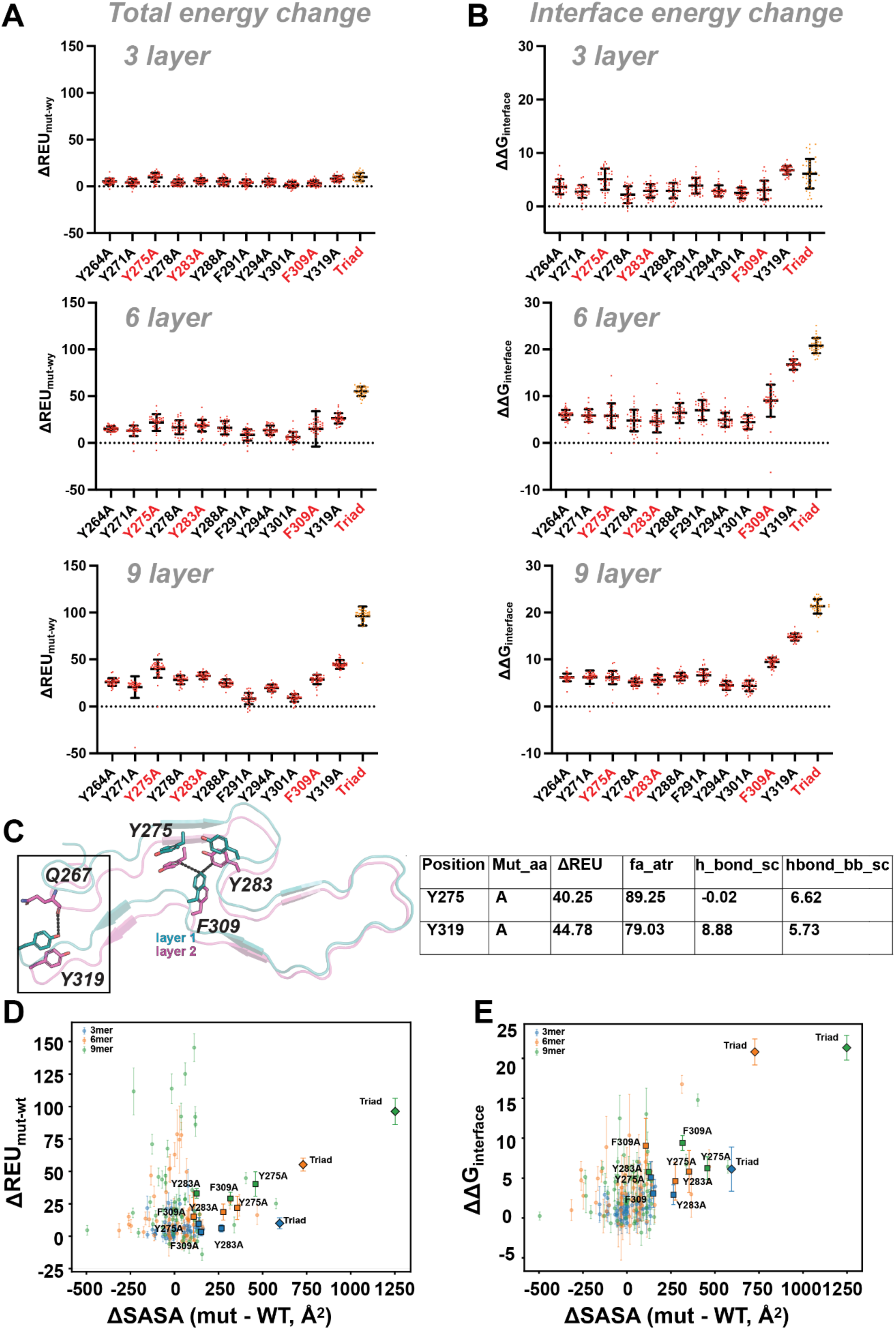
Aromatic substitutions and the Y275/Y283/F309 triad perturb hnRNPA2 fibril energetics and solvent exposure. **(A)** Total energetic changes (ΔREU_mut-WT_) for individual aromatic-to-alanine substitutions and the combined Y275A/Y283A/F309A triad in 3-, 6- and 9-layer fibril assemblies. Individual Rosetta replicates and summary values are shown as an average with standard deviation across 35 replicates. Y275A, Y283A, F309A and the triad are highlighted. **(B)** Corresponding ΔΔG_interface_ values for the same aromatic substitutions and triad across 3-, 6- and 9-layer assemblies. Individual Rosetta replicates and summary values are shown as an average with standard deviation across 35 replicates. **(C)** Structural view of selected aromatic interactions within adjacent fibril layers. Y275, Y283, F309 and Y319 are indicated, together with neighboring Q267. The accompanying table summarizes selected energetic terms for Y275A and Y319A. **(D)** Relationship between the change in total solvent-accessible surface area (ΔSASA, mutant − WT) and total energetic change for alanine substitutions in 3-, 6- and 9-layer assemblies. Y275A, Y283A, F309A and the triad are highlighted. Data are shown as average with standard deviation across 35 replicates for energetics and a single value for SASA (lowest energy structure). **(E)** Relationship between ΔSASA and ΔΔG_interface_. The Y275A/Y283A/F309A triad produces both a large increase in solvent exposure and a strong interface energetic penalty, particularly in the 6- and 9-layer assemblies. Data are shown as average with standard deviation across 35 replicates for energetics and a single value for SASA (lowest energy structure)

**SI Appendix, Figure 6.**
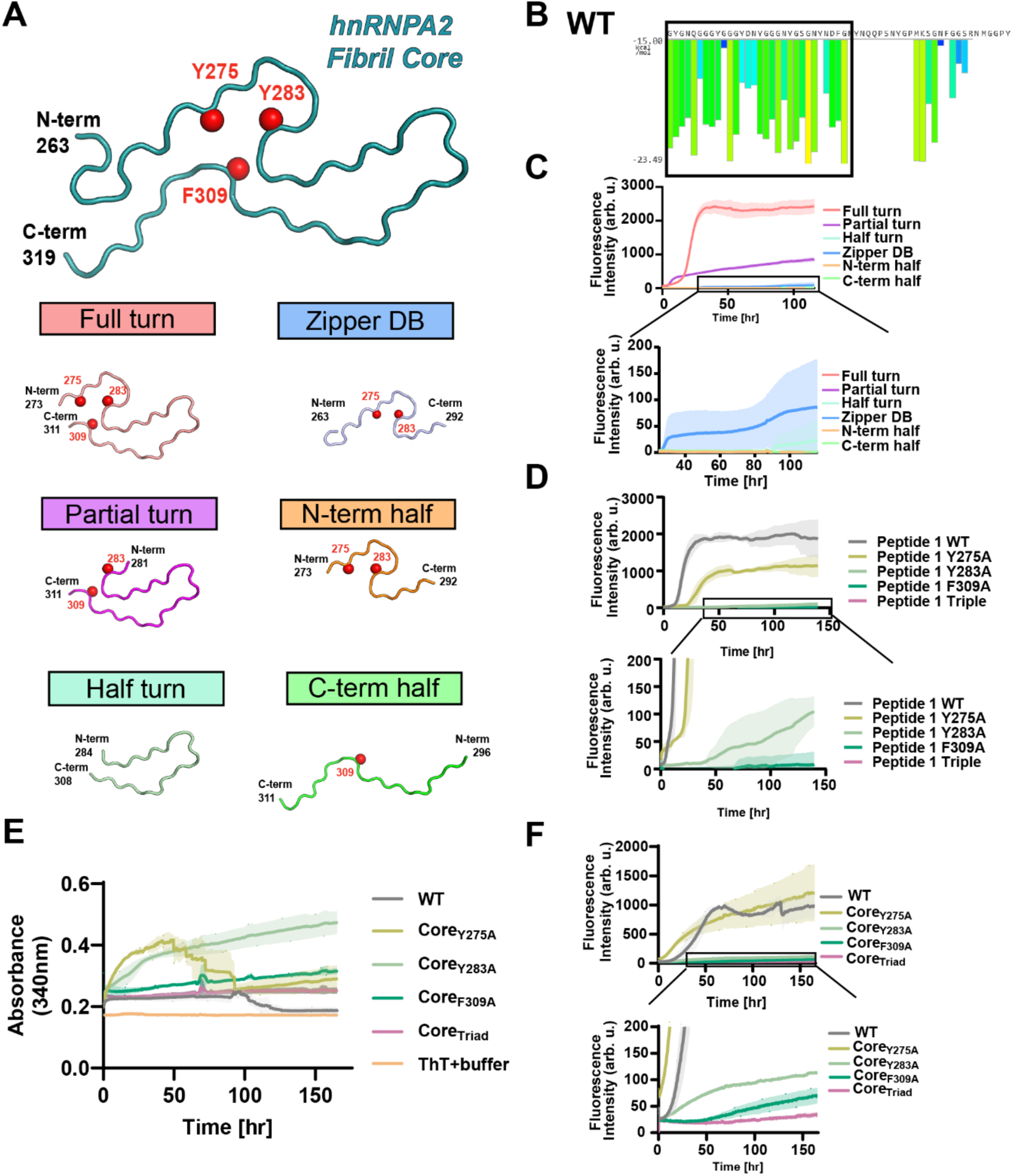
Sequence and aggregation analysis of hnRNPA2 fibril-core peptides and aromatic mutants. **(A)** Structural representation of the hnRNPA2 fibril core and peptide fragments used to define sequence requirements for fibril formation. Y275, Y283 and F309 are indicated within the fibril structure. Peptides correspond to the full turn, partial turn, half turn, ZipperDB-derived segment, N-terminal half and C-terminal half of the fibril core. **(B)** ZipperDB amyloid-propensity profiles for WT hnRNPA2. Boxed region represents the sequence of the ZipperDB-derived segment. **(C)** Representative ThT fluorescence traces for the indicated fibril-core peptides. The full-turn peptide undergoes robust aggregation, while the partial-turn construct retains aggregation activity with faster kinetics; the remaining fragments show little or no ThT increase under the conditions tested. Data are shown as an average across three replicates with standard deviation. **(D)** Representative ThT fluorescence traces for indicated full turn peptide mutants. Point mutants at F309 and Triad decrease aggregation propensity. Data are shown as an average across three replicates with standard deviation. **(E)** Turbidity measurements of mCherry-hnRNPA2-LCD mutants. All mutants have relatively low turbidity which then increases over time most likely due to fibril formation. **(F)** Representative ThT fluorescence traces for WT mCherry-hnRNPA2-LCD and the indicated core aromatic mutants. Core_Y275A_ retains substantial aggregation, whereas Core_Y283A_, Core_F309A_ and Core_Triad_ show progressively reduced ThT responses. Data are shown as an average across three replicates with standard deviation.

**SI Appendix, Figure 7.**
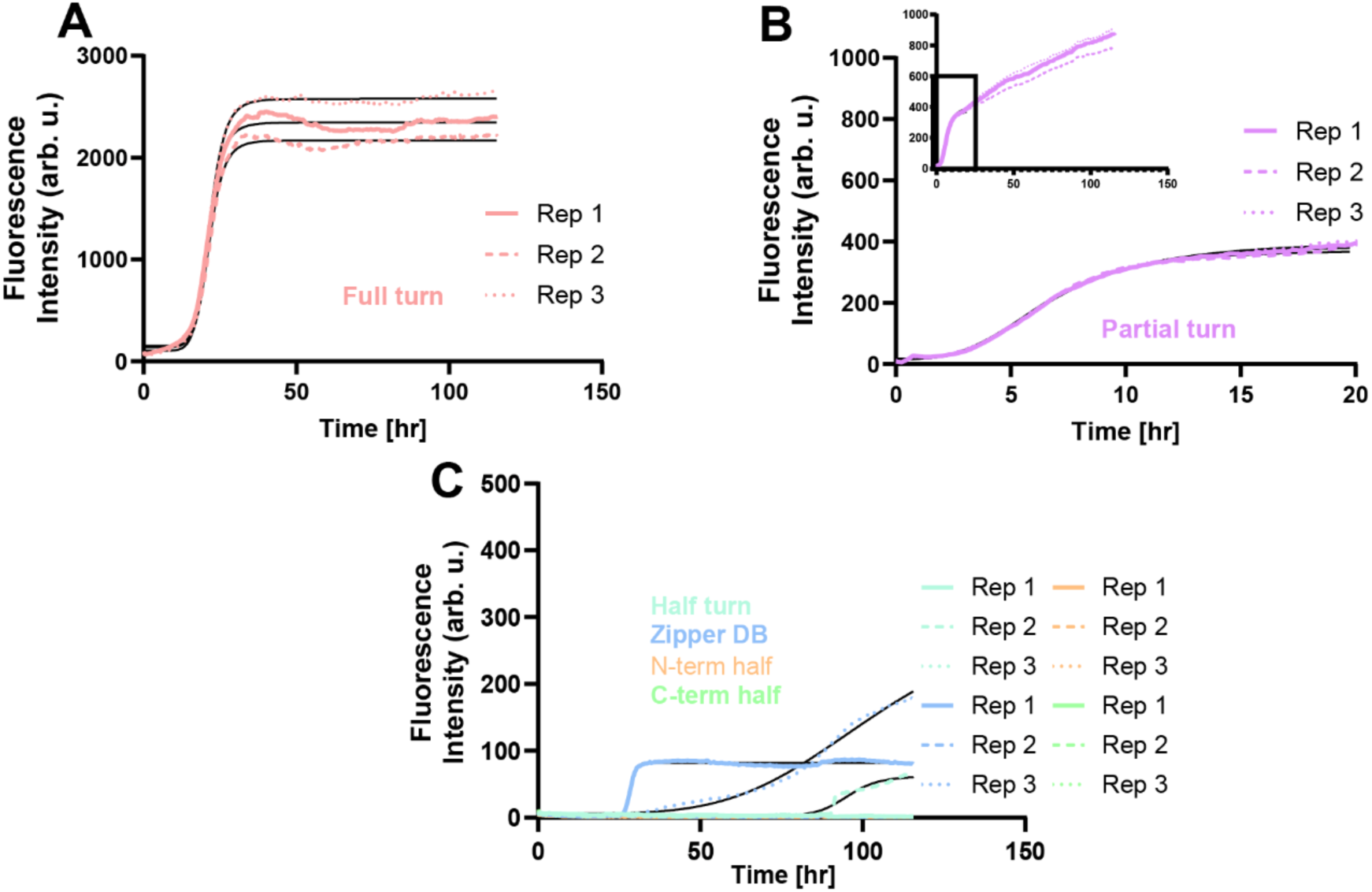
Raw ThT sigmoidal fits of full turn peptide point mutants. (A-C) Fits to the raw ThT fluorescence curves were used to estimate T_1/2_ values. Zoomed in ThT curves are shown when the curve was truncated to achieve a more accurate fit. Data are shown as individual replicates with overlaid fits.

**SI Appendix, Figure 8.**
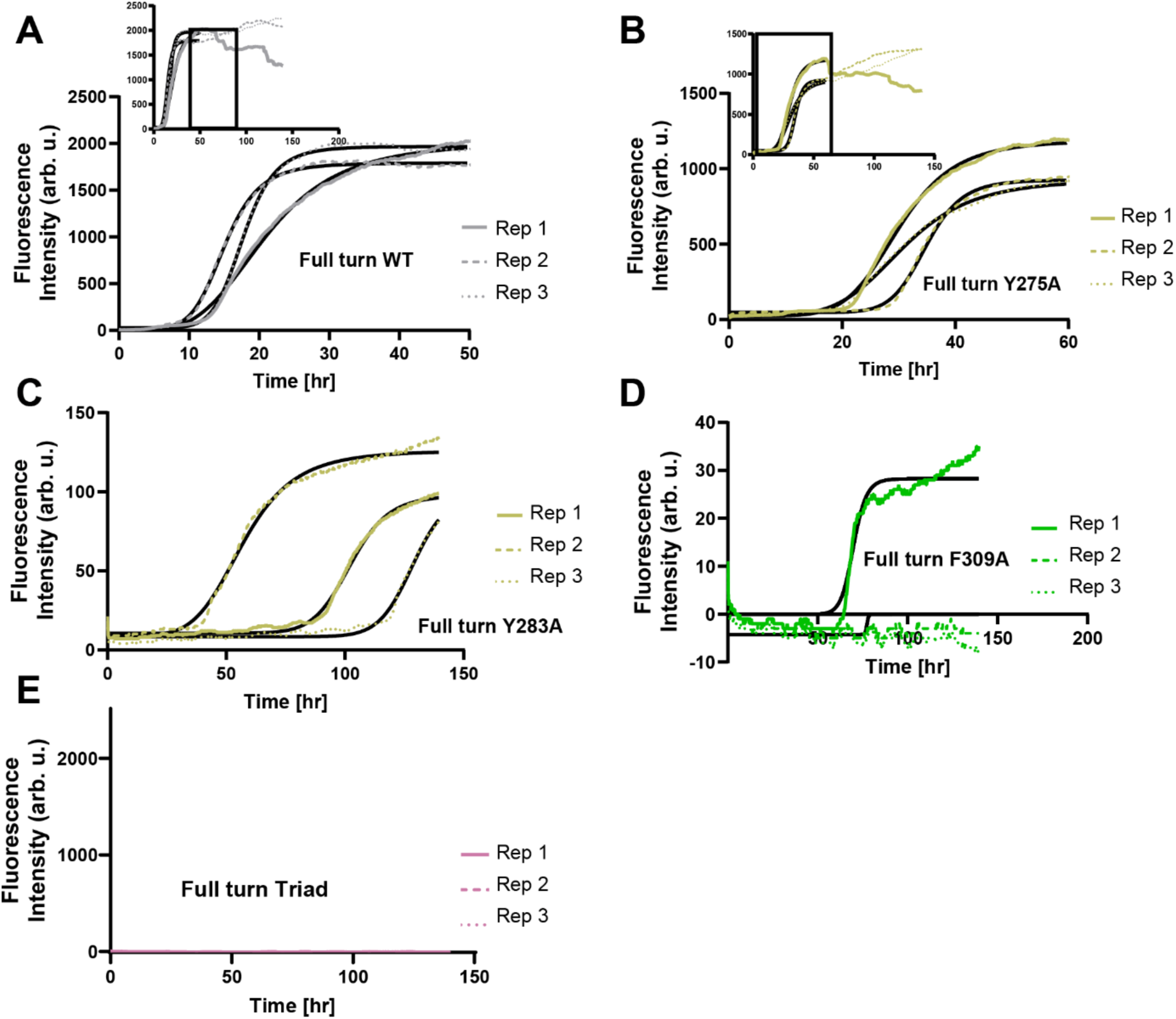
Raw ThT sigmoidal fits of full turn peptide point mutants. (A-E) Fits to the raw ThT fluorescence curves were used to estimate T_1/2_ values. Zoomed in ThT curves are shown when the curve was truncated to achieve a more accurate fit. Data are shown as individual replicates with overlaid fits.

**SI Appendix, Figure 9.**
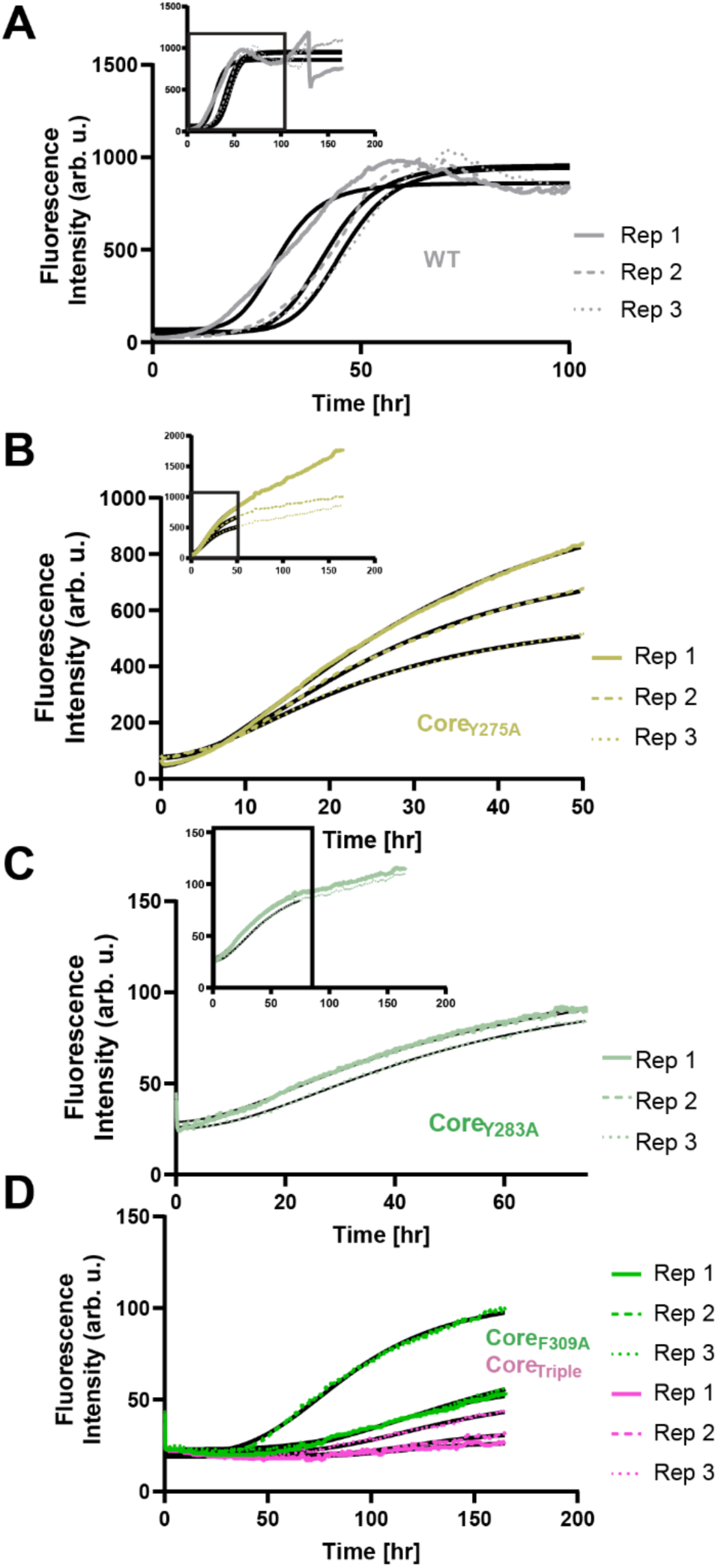
Raw ThT sigmoidal fits of mCherry-hnRNPA2 constructs. (A-D) Fits to the raw ThT fluorescence curves were used to estimate T_1/2_ values. Zoomed in ThT curves are shown when the curve was truncated to achieve a more accurate fit. Data are shown as individual replicates with overlaid fits.

**SI Appendix, Figure 10.**
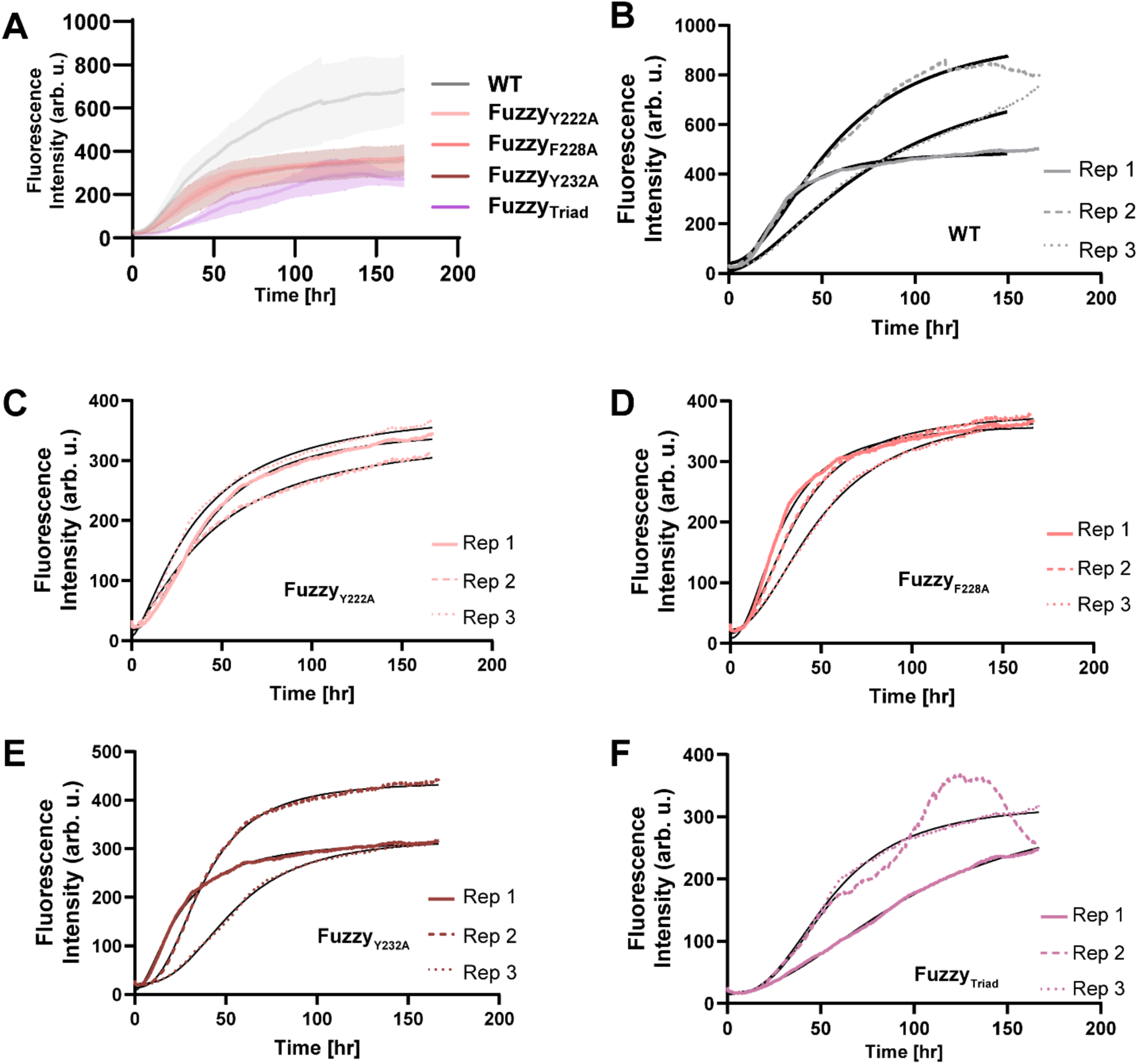
Fuzzy coat mutants in the mCherry-hnRNPA2 LCD construct do not inhibit aggregation in the one-phase regime. (A) Raw ThT curves for each mutant. Data are shown as averages from three replicates with standard deviation. (B-F) Fits to the raw ThT fluorescence curves for WT and each mutant were used to estimate T_1/2_ values across replicate experiments. Data are shown as individual replicates with overlaid fits.

**SI Appendix, Figure 11.**
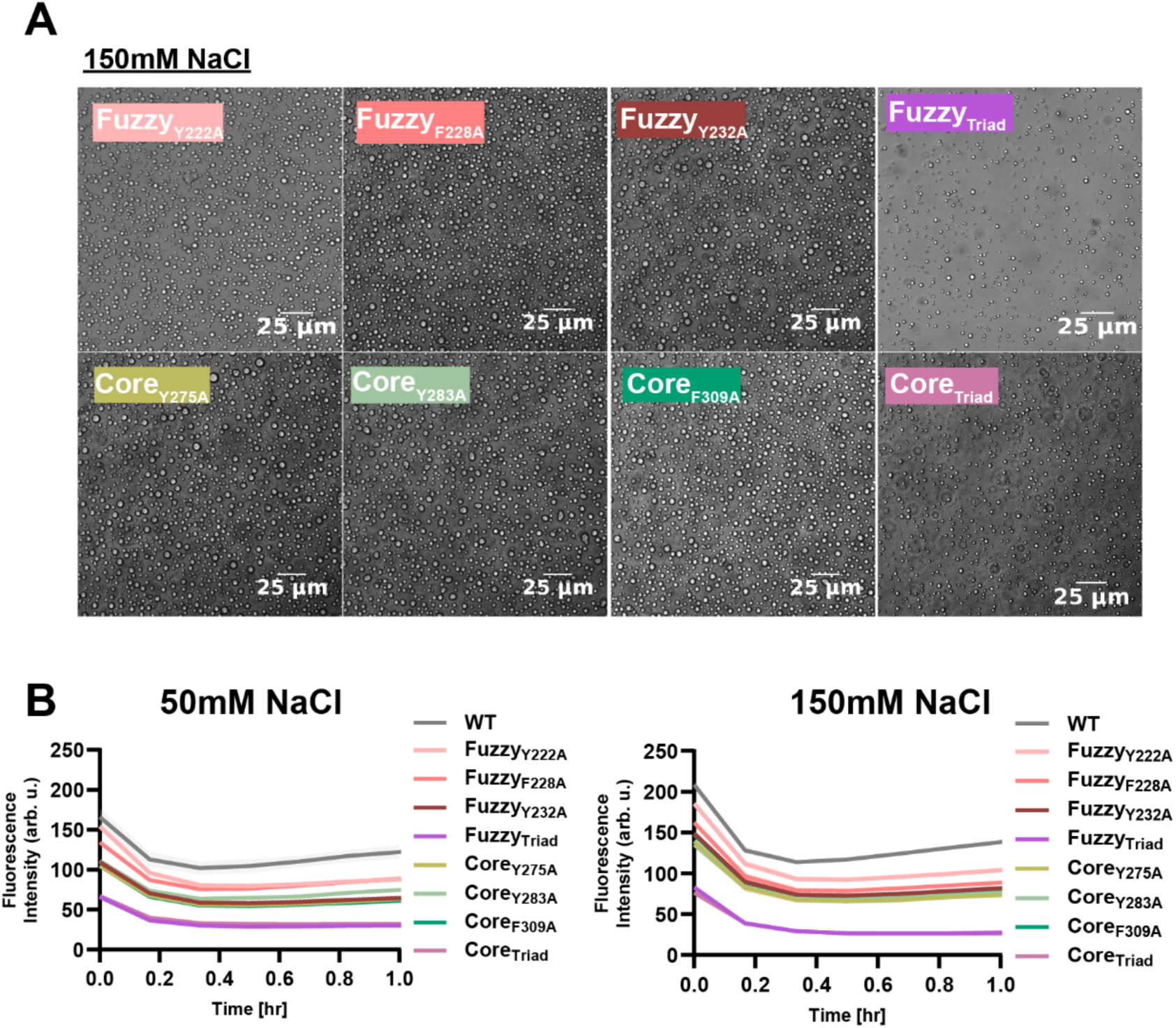
Micrographs of condensates induced at 150 mM NaCl and ThT curves at 1hr. **(A)** Representative light-microscopy images of condensates formed by the indicated fuzzy-coat and fibril-core aromatic mutants at 150 mM _NaCl_. Scale bars, 25 μm. **(B)** ThT curves for all replicate 2 hnRNPA2 mutants in the two-phase regime, truncated at one hour showing no increase in ThT signal during this timeframe. Data are shown as an average across three replicates.

**SI Appendix, Figure 12.**
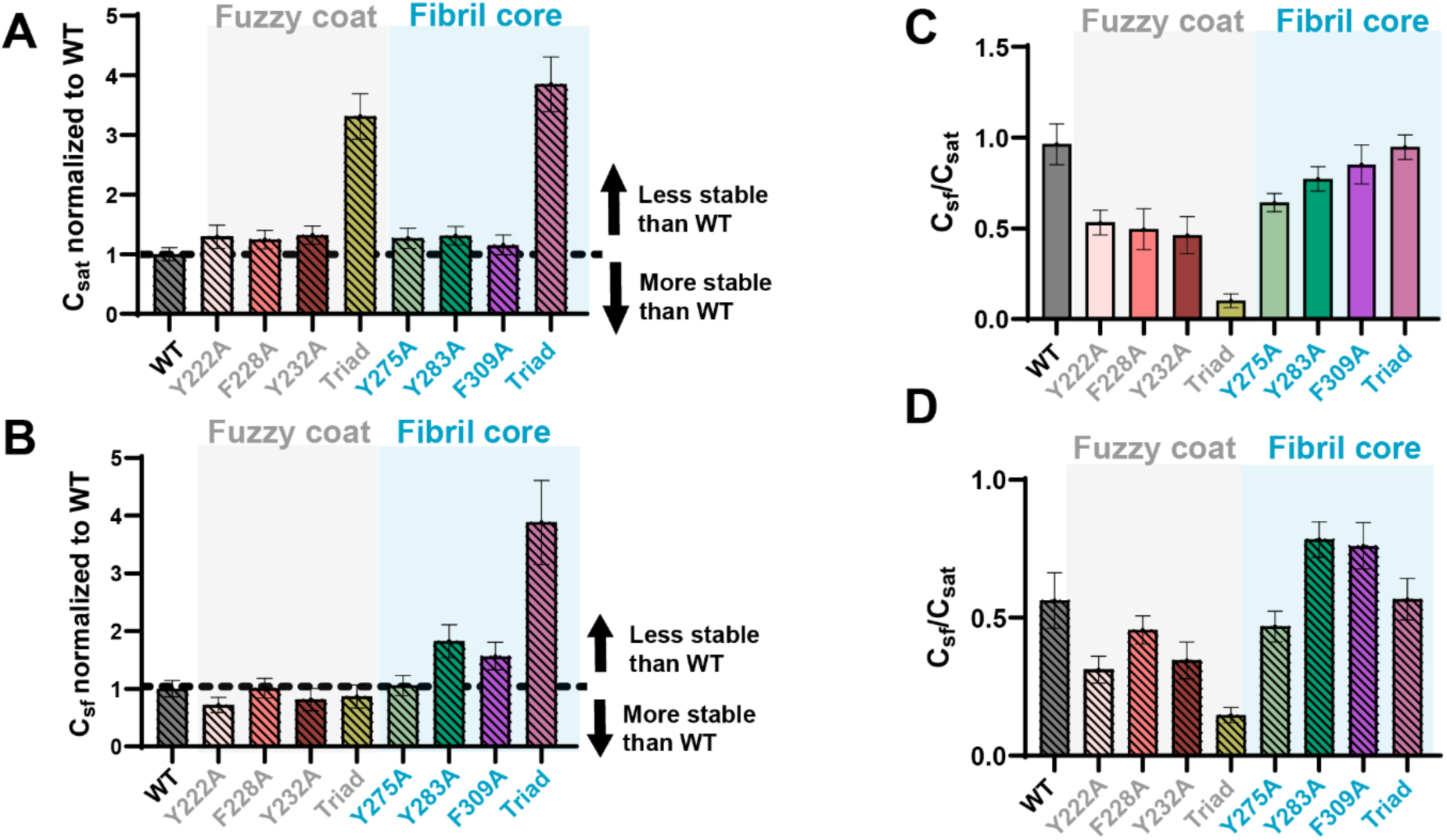
C_sat_ and C_sf_ values at 150 mM NaCl and C_sf_/C_sat_ ratios for all constructs. (A-B) C_sat_ and C_sf_ values normalized to WT, for constructs incubated at 150 mM NaCl. Values above 1 indicate condensates/fibrils are less stable, whereas values below 1 indicate condensates/fibrils are more stable. Fuzzy_Triad_ and Core_Triad_ similarly increase C_sat_, whereas Core_Triad_ produces the largest increase in C_sf_. Error bars represent SEM. **(C-D)** C_sf_/C_sat_ ratios for WT and all constructs. The largest separation between fuzzy-coat and core variants is observed for the triad mutants. Error bars represent SEM.

**SI Appendix, Figure 13.**
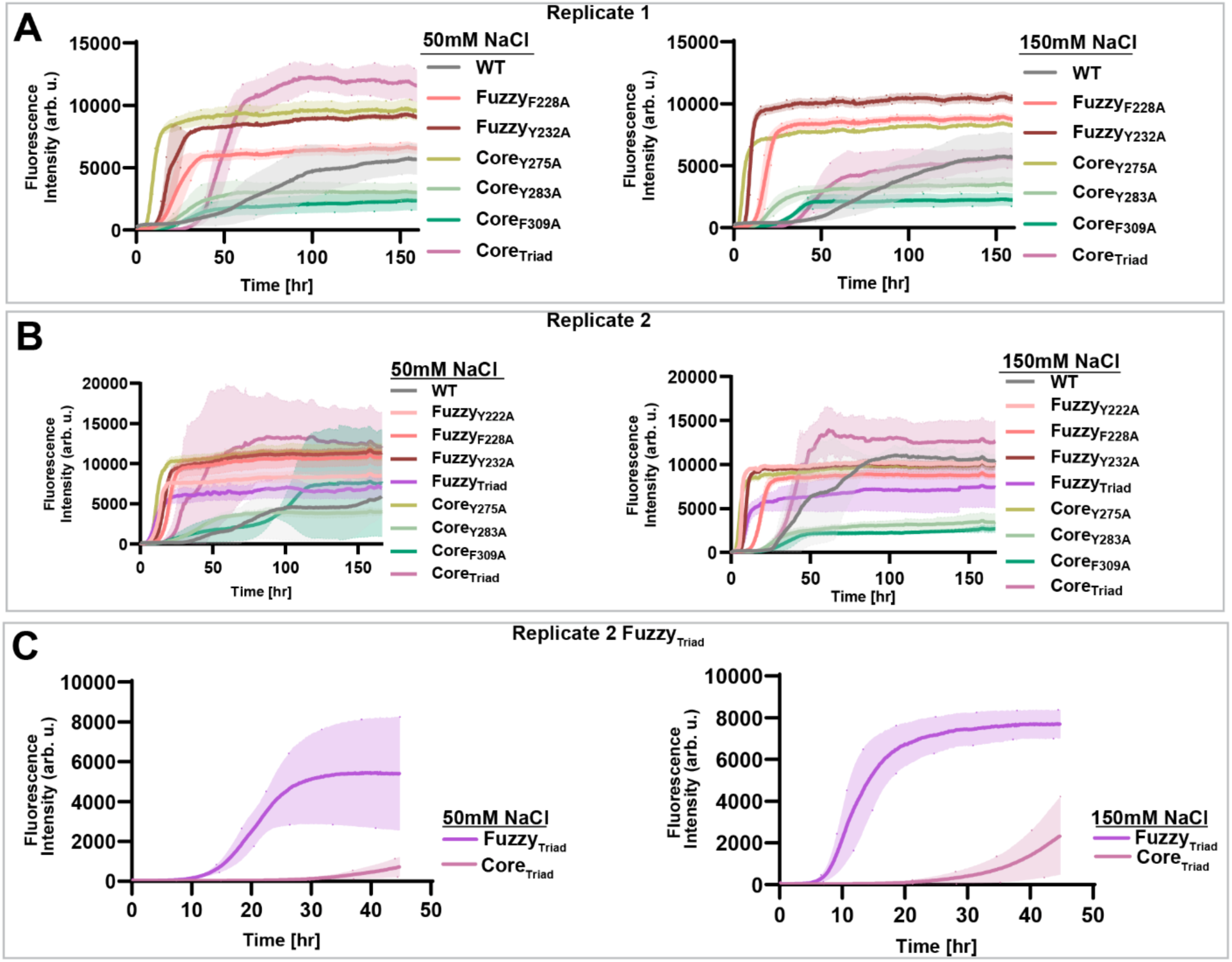
Replicate ThT measurements of hnRNPA2 fibril formation in the two-phase regime. **(A)** ThT fluorescence kinetics for WT and the indicated fuzzy- coat and fibril-core aromatic mutants following induction of phase separation at 50 or 150 mM NaCl in replicate 1. Data are shown as averages with shaded region representing SD. **(B)** Independent replicate (replicate 2) ThT measurements at 50 and 150 mM NaCl. Shaded regions indicate SD associated with the kinetic measurements. Data are shown as averages with shaded region representing SD. **(C)** Independent replicate of Fuzzy_Triad_ and CoreTriad aggregation kinetics at 50 and 150 mM NaCl. Fuzzy_Triad_ undergoes substantially faster ThT-positive conversion than Core_Triad_ despite both mutations disrupting three aromatic residues. Data are shown as averages from three technical replicates with SD.

**SI Appendix, Figure 14.**
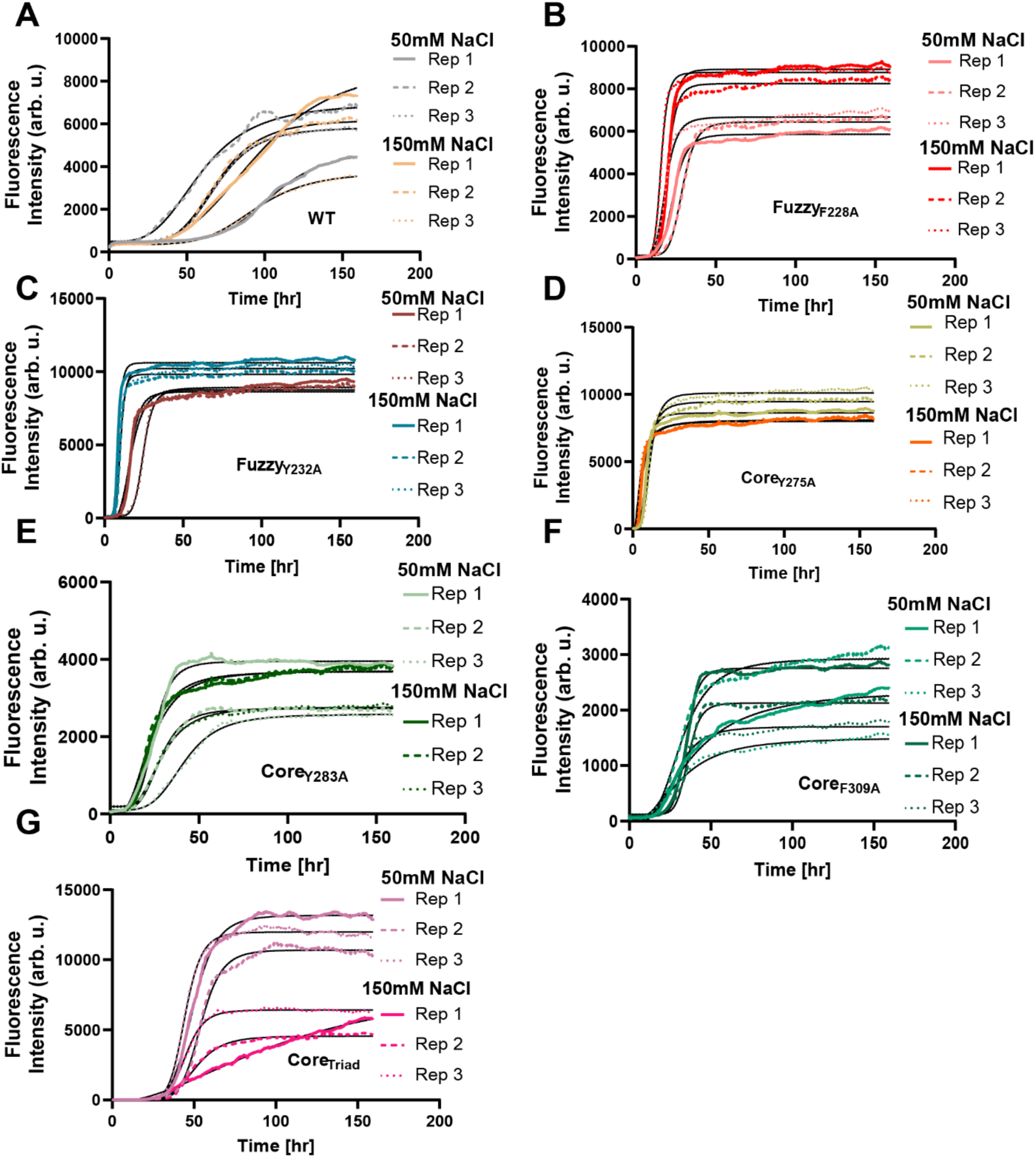
Raw ThT sigmoidal fits of untagged hnRNPA2 constructs (replicate 1). (A-G) Fits to the raw ThT fluorescence curves for WT and mutants were used to estimate T_1/2_ values across replicate experiments. Data are shown as individual replicates overlaid with fitted curves.

**SI Appendix, Figure 15.**
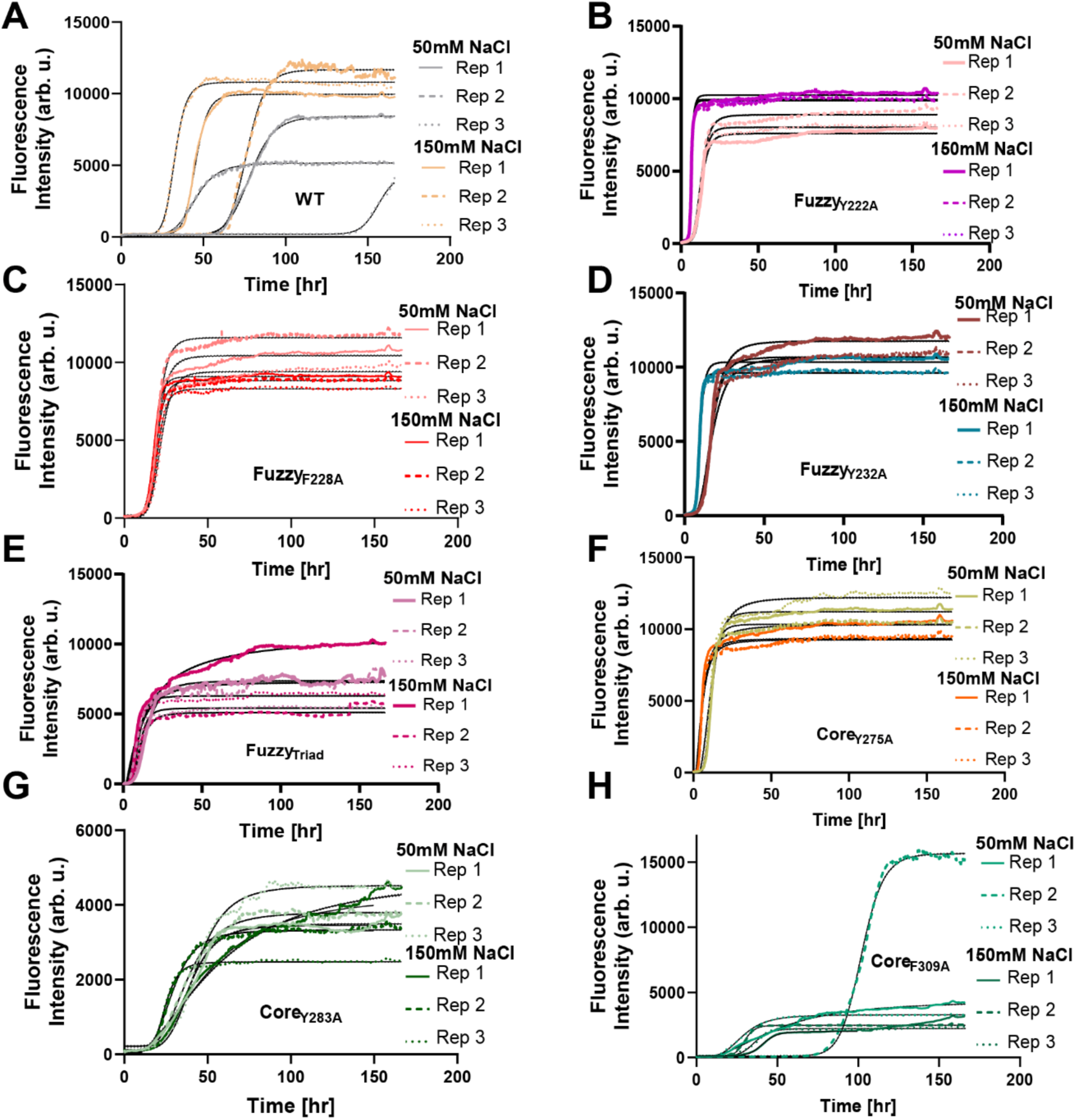
Raw ThT sigmoidal fits of untagged hnRNPA2 constructs (replicate 2). (A-H) Fits to the raw ThT fluorescence curves for WT and mutants were used to estimate T_1/2_ values across replicate experiments. Data are shown as individual replicates overlaid with fitted curves.

**SI Appendix, Figure 16.**
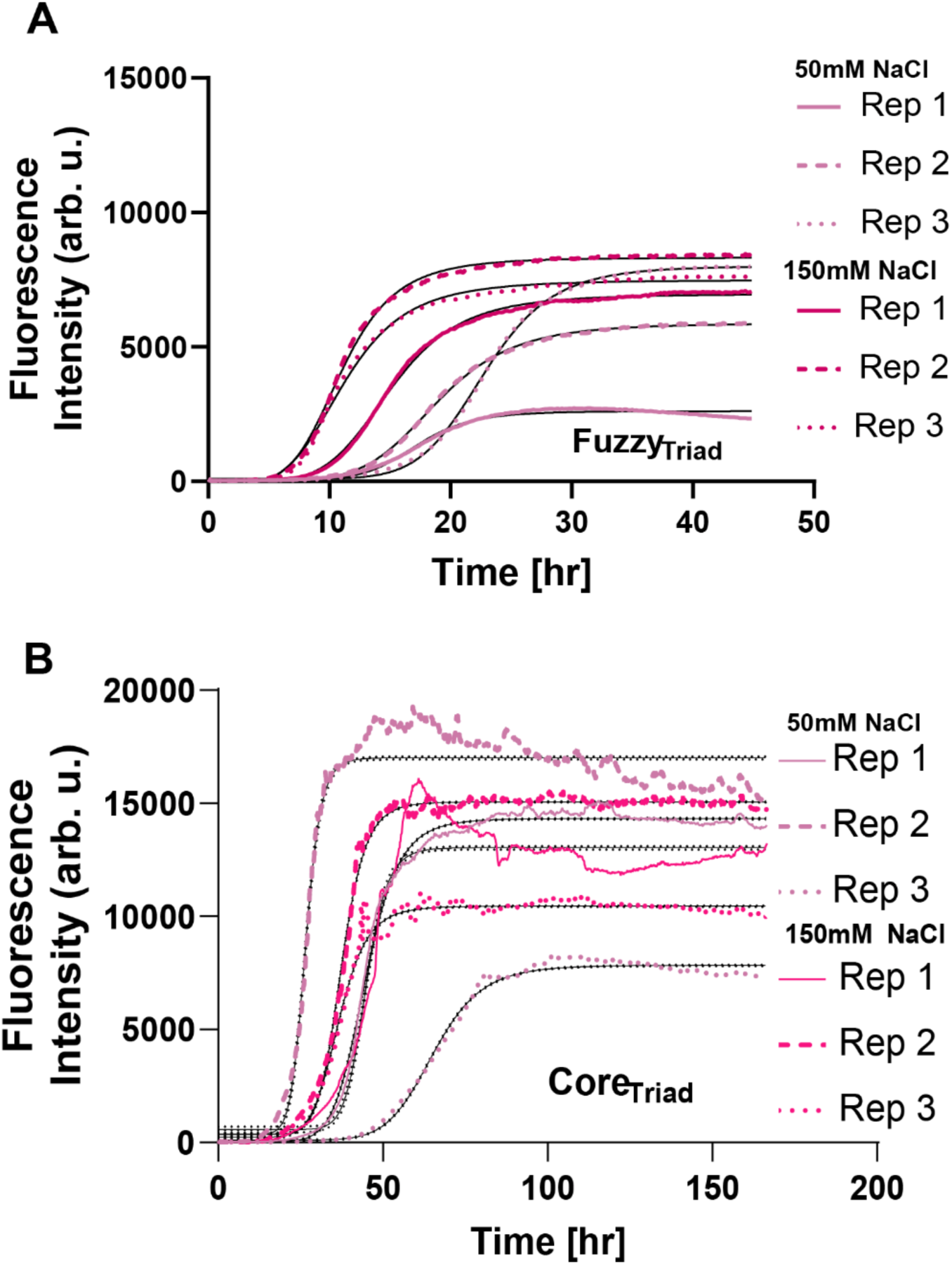
Raw ThT sigmoidal fits of untagged hnRNPA2 constructs (replicate 2 cont. and independent replicate with Fuzzy_Triad_). (A-B) Fits to the raw ThT fluorescence curves for each triad mutant were used to estimate T_1/2_ values across replicate experiments. Data are shown as individual replicates overlaid with fitted curves.

**SI Appendix, Figure 17.**
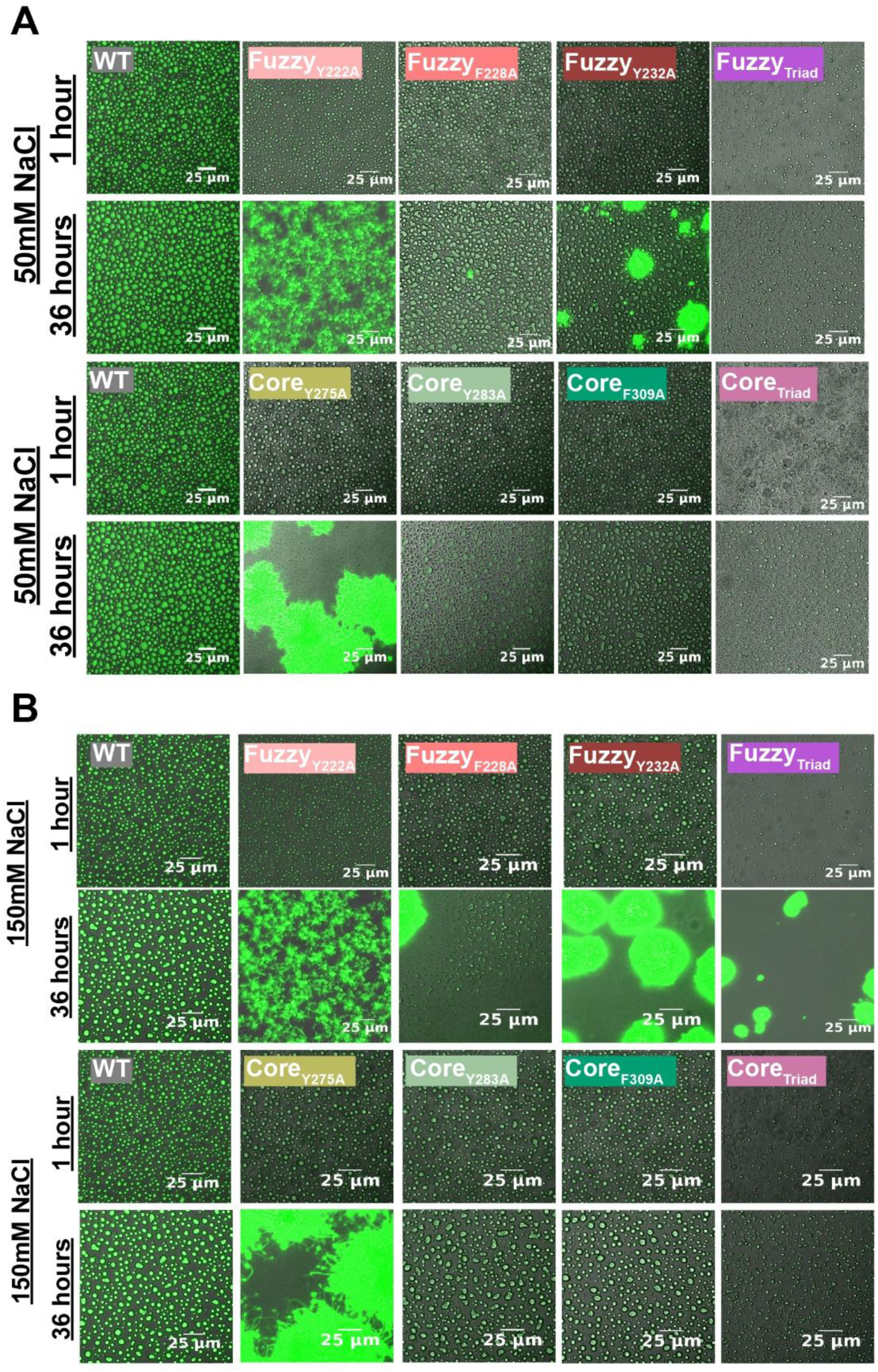
Time-dependent condensate-to-aggregate conversion of hnRNPA2 aromatic mutants at 50 mM NaCl and 150 mM NaCl. (A-B) Representative microscopy images of WT hnRNPA2-LCD and fuzzy-coat or fibril core aromatic mutants at 1 and 36 h after induction of phase separation at 50 mM NaCl (A) and 150 mM NaCl (B). WT retains abundant droplet-like structures over the imaging period, whereas several aromatic mutants undergo pronounced time-dependent changes in morphology. Fuzzy-coat mutants and Core_Y275A_ show particularly rapid formation of larger aggregated structures, whereas Core_Y283A_, Core_F309A_ and Core_Triad_ retain more droplet-like morphologies. Scale bars, 25 μm.

**SI Appendix, Figure 18.**
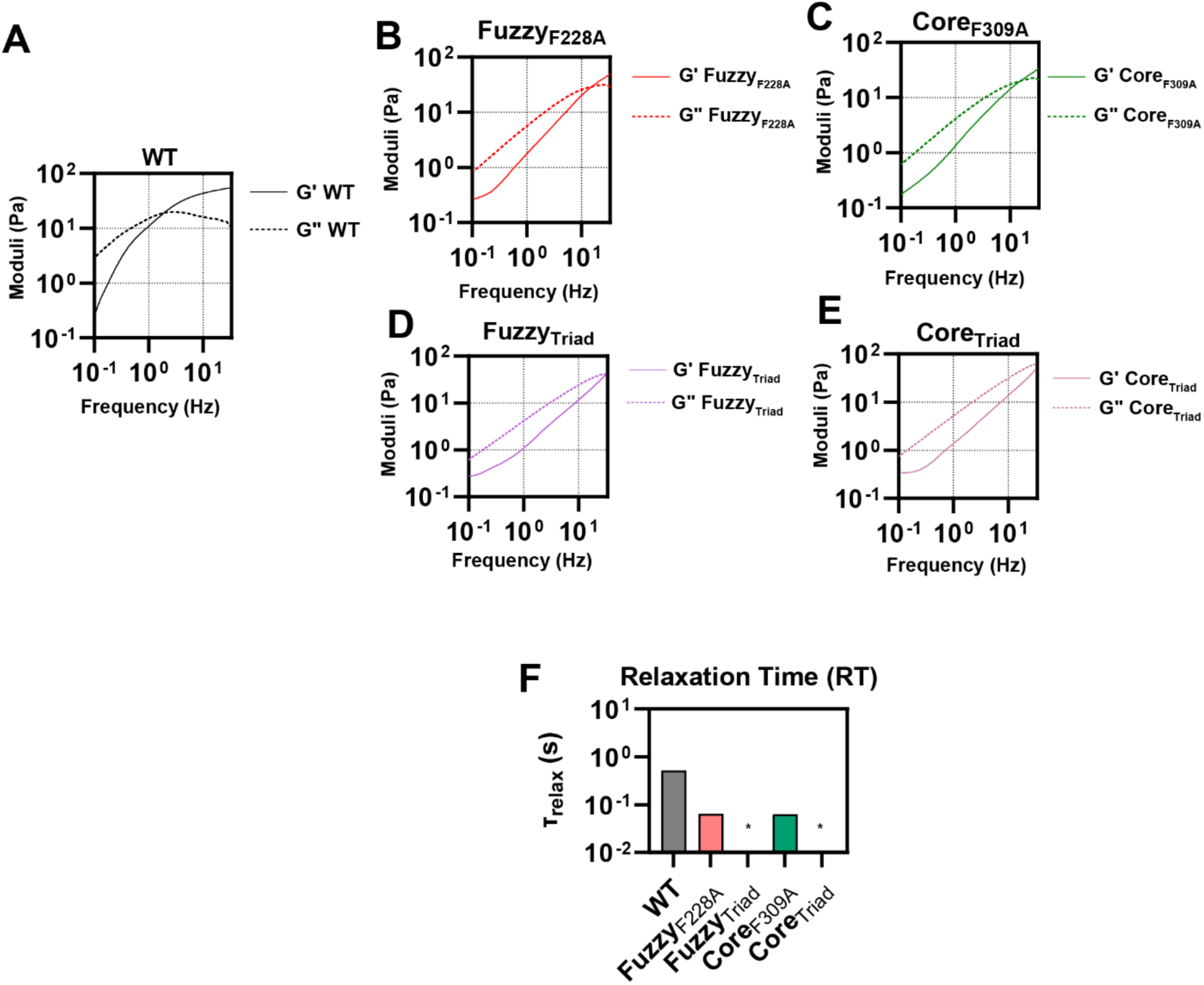
Frequency-dependent viscoelastic properties and relaxation times of hnRNPA2 condensates. Frequency-dependent storage (G’, solid lines) and loss (G’’, dashed lines) moduli derived from video particle-tracking microrheology for **(A)** WT, **(B)** Fuzzy_F228A_, **(C)** Core_F309A_, **(D)** Fuzzy_Triad_ and **(E)** Core_Triad_ hnRNPA2-LCD condensates. At low frequencies, the loss modulus dominates, consistent with terminally viscous behavior, whereas the elastic contribution increases with frequency. **(F)** Terminal relaxation times (τ_relax_) calculated from the crossover between G’ and G’’. WT condensates exhibit a substantially longer relaxation time than the single aromatic mutants. The crossover frequencies for Fuzzy_Triad_ and Core_Triad_ fall outside the experimentally accessible range, indicating relaxation on timescales shorter than those resolved by the measurement.

## SI Appendix, Movie legends

Movie S1 (separate file): 24hr time lapse of hnRNPA2-LCD WT condensates in 50 mM NaCl

Movie S2 (separate file): 24hr time lapse of hnRNPA2-LCD Fuzzy_Y222A_ condensates in 50 mM NaCl

Movie S3 (separate file): 24hr time lapse of hnRNPA2-LCD Fuzzy_F228A_ condensates in 50 mM NaCl

Movie S4 (separate file): 24hr time lapse of hnRNPA2-LCD Fuzzy_Y232A_ condensates in 50 mM NaCl

Movie S5 (separate file): 24hr time lapse of hnRNPA2-LCD Core_Y275A_ condensates in 50 mM NaCl

Movie S6 (separate file): 24hr time lapse of hnRNPA2-LCD Core_Y283A_ condensates in 50 mM NaCl

Movie S7 (separate file): 24hr time lapse of hnRNPA2-LCD Core_F309A_ condensates in 50 mM NaCl

Movie S8 (separate file): 24hr time lapse of hnRNPA2-LCD Core_Triad_ condensates in 50 mM NaCl

Movie S9 (separate file): 24hr time lapse of hnRNPA2-LCD Fuzzy_Triad_ condensates in 50 mM NaCl

Movie S10 (separate file): 24hr time lapse of hnRNPA2-LCD WT condensates in 150 mM _NaCl_

Movie S11 (separate file): 24hr time lapse of hnRNPA2-LCD Fuzzy_Y222A_ condensates in 150 mM NaCl

Movie S12 (separate file): 24hr time lapse of hnRNPA2-LCD Fuzzy_F228A_ condensates in 150 mM NaCl

Movie S13 (separate file): 24hr time lapse of hnRNPA2-LCD Fuzzy_Y232A_ condensates in 150 mM NaCl

Movie S14 (separate file): 24hr time lapse of hnRNPA2-LCD Core_Y275A_ condensates in 150 mM NaCl

Movie S15 (separate file): 24hr time lapse of hnRNPA2-LCD Core_Y283A_ condensates in 150 mM NaCl

Movie S16 (separate file): 24hr time lapse of hnRNPA2-LCD Core_F309A_ condensates in 150 mM NaCl

Movie S17 (separate file): 24hr time lapse of hnRNPA2-LCD Core_Triad_ condensates in 150 mM NaCl

Movie S18 (separate file): 24hr time lapse of hnRNPA2-LCD Fuzzy_Triad_ condensates in 150 mM NaCl

## Notes

### Competing Interest Statement

The authors have declared no competing interest.

